# Nutrient and salt depletion synergistically boosts glucose metabolism in individual bacteria

**DOI:** 10.1101/2022.01.26.477826

**Authors:** Georgina Glover, Margaritis Voliotis, Urszula Łapińska, Brandon M. Invergo, Darren Soanes, Paul O’Neill, Karen Moore, Nela Nikolic, Peter G. Petrov, David S. Milner, Sumita Roy, Kate Heesom, Thomas A. Richards, Krasimira Tsaneva-Atanasova, Stefano Pagliara

## Abstract

The interaction between a cell and its environment shapes fundamental intracellular processes such as cellular metabolism. In most cases growth rate is treated as a proximal metric for understanding the cellular metabolic status. However, changes in growth rate might not reflect metabolic variations in individuals responding to environmental fluctuations. Here we use single-cell microfluidics-microscopy combined with transcriptomics, proteomics and mathematical modelling to quantify the accumulation of glucose within *Escherichia coli* cells. In contrast to the current consensus, we reveal that environmental conditions which are comparatively unfavourable for growth, where both nutrients and salinity are depleted, increase glucose accumulation rates in individual bacteria and population subsets. We find that these changes in metabolic function are underpinned by variations at the translational and posttranslational level but not at the transcriptional level and are not dictated by changes in cell size. The metabolic response-characteristics identified greatly advance our fundamental understanding of the interactions between bacteria and their environment and have important ramifications when investigating cellular processes where salinity plays an important role.

## INTRODUCTION

All core metabolic networks require the uptake and utilization of carbon sources. Such function is variant, adaptable and in the process of being precisely shaped by natural selection as many heterotrophic organisms need to adapt their metabolic capabilities to dwell in scarce nutrient environments[1–4]. In the case of bacteria, sugars often represent the primary driving force for growth; they are used to power replication, to make storage compounds, and/or for the production of secondary metabolites that further dictate metabolic function[5, 6]. Glucose is commonly employed to investigate the regulation of sugar uptake and metabolism in bacteria[7]. In fact, many bacterial species primarily use glucose when exposed to nutrient mixtures[8, 9] and have evolved several independent ways of acquiring glucose from the environment, as characterised by hundreds of variant glucose transport systems[8].

In gram-negative bacteria, such as *Escherichia coli*, glucose passively diffuses through outer membrane porins whose expression is regulated both at the transcriptional and translational level[10–12]. Glucose then crosses the *E. coli* inner membrane via five different permeases including the glucose and mannose phosphotransferase systems (PTS)[8],[13]. Once in the cytoplasm glucose is phosphorylated to glucose-6-phosphate that is broken down to pyruvate, this in turn is metabolised to acetyl-CoA which then enters the citric acid cycle[14] generating ATP. At micromolar extracellular concentrations, glucose and its fluorescent analogue 2-(N-(7-Nitrobenz-2-oxa-1,3-diazol-4-yl)Amino)-2-Deoxyglucose (2-NBDG) transiently accumulate intracellularly as a result of concomitant uptake and degradation processes^15^. Seminal experiments carried out in chemostats suggest that glucose uptake increases with cell surface area[5,15–17] and it is heterogeneous within clonal *E. coli* populations.

These insights were obtained by employing bacteria growing in optimal conditions or at low nutrient levels[15]. In contrast, in natural environments changes in other parameters are common place[18–24]. For example, the salinity of aquatic ecosystems is changing worldwide[25], it has been rated as one of the most important environmental changes[26] and can have a profound impact on microbial growth and infectivity[27, 28], thus affecting biodiversity[29]. Salinity is decreasing in naturally saline ecosystems due to increased agricultural drainages and unusual wind patterns[30], whereas the salinity of freshwater ecosystems is increasing due to de-icing or salt mining (up to 97% increase[29]), along with climatic aridification and rising sea levels[25]. Furthermore, salinity can dramatically change across different bodily environments with a ∼7-fold decrease in NaCl content from the colon to the ileum (i.e from 6.1 to 0.9 g/L)[31–33] or across different humans with a 40% increase in the lungs of cystic fibrosis patients[34]. These salinity variations often occur concomitantly other environmental variations, such as changes in temperature, pH or nutrient levels, which can result in additive, antagonistic or synergistic effects on microbes including *Escherichia coli*[35, 36]. Variations in temperature or content of metals have frequently been investigated in combination with salinity, whereas changes in the nutrient-salinity pair has received less attention[35]. Understanding and predicting the effects of multiple environmental variations on the phenotypic diversity in microbial traits, such as metabolic rates, is critical for unravelling how microbes function in their environment.

Here we study the diversity in glucose metabolism within clonal *E. coli* populations and show that simultaneous extracellular nutrient and salt depletion synergistically enhance glucose metabolism. These single-cell traits are not displayed when bacteria are exposed to nutritional or salinity depletion alone, demonstrating that the effect of these environmental changes on glucose metabolism is not additive. These changes in metabolic function are underpinned by variations at the translational and posttranslational level but not at the transcriptional level and are not dictated by changes in cell size. These findings offer unique understanding of the interaction between a cell and its environment and will inform modelling metabolic flux across bacterial populations[6], adjusting process parameters in biotechnology, food preservation or metabolic engineering[37] and optimizing treatment with antibiotics that utilize sugar uptake pathways to reach their intracellular target[38–40].

## RESULTS

### Experimental assessment of glucose accumulation and degradation in individual bacteria

In order to quantify glucose uptake and accumulation in individual bacteria, we introduced a clonal *E. coli* population into a microfluidic mother machine device[41]. This device is equipped with a large microfluidic chamber for bacteria loading and media delivery via pressure-driven microfluidics, connected with thousands of bacteria hosting channels with cross section comparable to individual bacteria (inset in Fig. 1).

**Figure 1.**
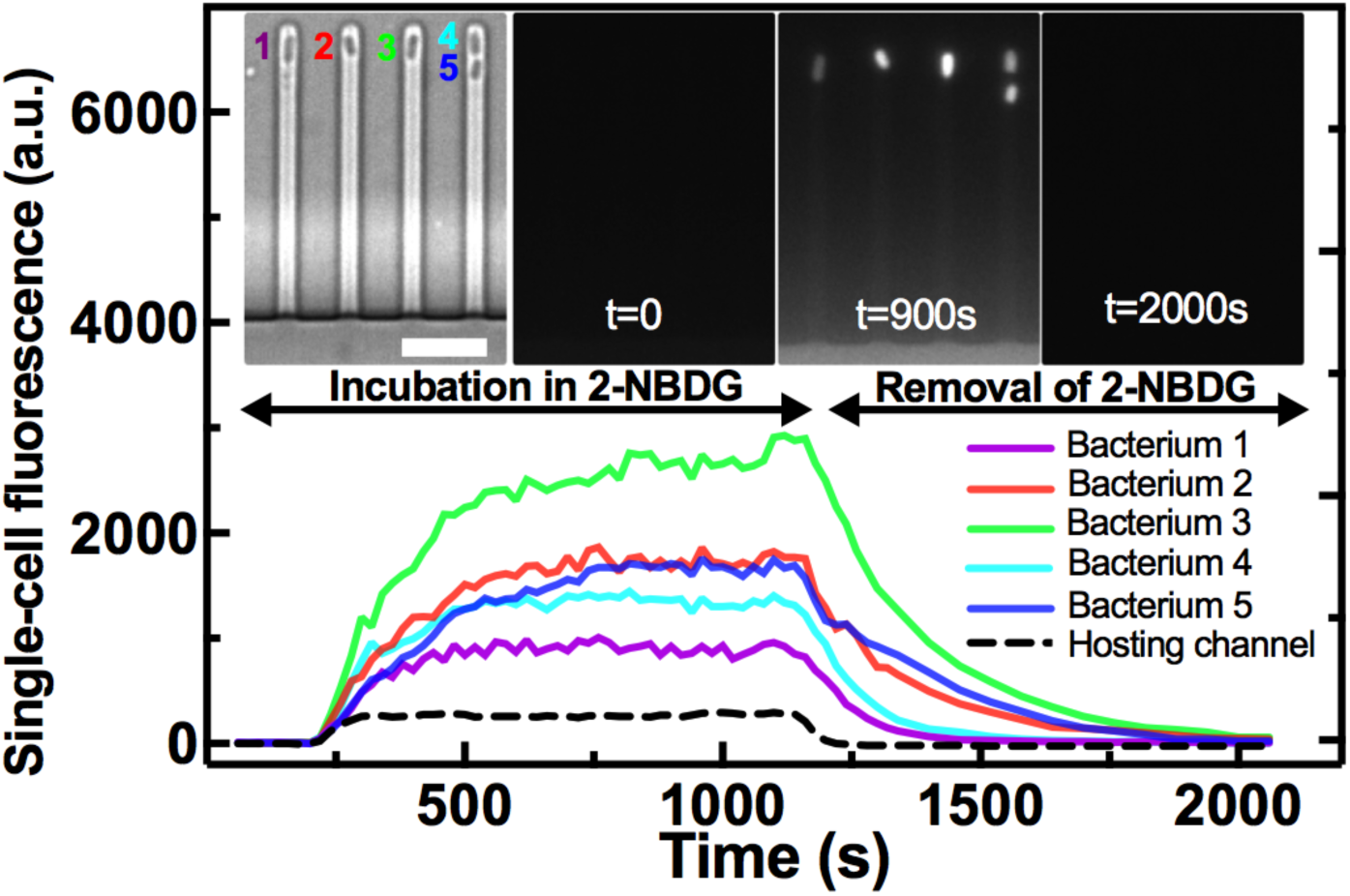
Experimental assessment of glucose accumulation and degradation in individual bacteria. Temporal dependence of the intracellular fluorescence of the glucose analogue 2-NBDG in five representative *E. coli* cells confined in the bacterial hosting channels of the microfluidic mother machine. The dashed line represents the fluorescence of the extracellular 2-NBDG in the bacterial hosting channels. Noteworthy, we measured 2-NBDG fluorescence as the mean fluorescent values of each pixel constituting each bacterium, thus normalizing by cell size. Insets from left to right: brightfield image of the five bacteria at *t*=0 s when the fluorescent glucose analogue is added to the mother machine; corresponding image in the green fluorescent channel; saturation of 2-NBDG intracellular fluorescence levels at *t*=900 s followed by removal of extracellular 2-NBDG at *t*=1200 s; complete degradation of intracellular 2-NBDG leading to intracellular fluorescence down to background levels by *t*=2000 s. Scale bar: 5 µm.

We then added 30 µM 2-NBDG (i.e. a fluorescent glucose analogue) dissolved in glucose-free M9 medium[5, 15] in the microfluidic chamber while measuring its diffusion in the bacteria hosting channels over time (dashed line in Fig. 1). We measured a progressive increase in the fluorescence of individual bacteria that quickly became brighter than the channel fluorescence (solid lines in Fig. 1 and insets) demonstrating higher intracellular compared to extracellular 2-NBDG concentration. This was due to the uptake of 2-NBDG and subsequent accumulation in single bacteria up to a steady-state[5,15,42,43]. This hyperbolic accumulation kinetics indicated the presence of a sink term, namely intracellular phosphorylation of 2-NBDG and its degradation by *E. coli*, processes that are comparable between glucose and 2-NBDG[5, 42].

To gain further insight of the dynamics of this degradation process, the extracellular 2-NBDG was washed away from the microfluidic chamber at *t*=1200 s and replaced with fresh LB medium. Consequently, we measured an exponential decrease in the fluorescence of both the bacteria hosting channels and of each bacterium (dashed and solid lines, respectively, in Fig. 1 and insets). This rapid decrease could not be accounted for by dilution due to cell growth[5, 44].

In order to quantify phenotypic heterogeneity in glucose accumulation, we evaluated the coefficient of variation (CV, the ratio between the standard deviation and the mean) of single-bacterium 2-NBDG fluorescence values across the clonal population at each time point. We verified that this heterogeneity could not be attributed to anisotropy in 2-NBDG concentration within the bacteria hosting channels (Fig. S1 and Table S1)[41].

### Glucose accumulation is maximal under simultaneous nutritional and salinity depletion

We then used the experimental approach above to determine the impact of nutritional and salinity depletion on glucose accumulation. We pre-cultured *E. coli* in three environments with three different salt contents broadly recapitulating the salinity encountered by bacteria in mesohaline (10 g/L NaCl), oligohaline (5 g/L NaCl) or fresh (0.5 g/L NaCl) water[29]. These salinity variations also approximate the NaCl concentration faced by bacteria in the colonic or the ileal environment (6.1 and 0.9 g/L NaCl, respectively)[32]. Moreover, at the lowest salinity, ion availability can become a rate-limiting factor[45, 46]. In each environment, *E. coli* were firstly pre-cultured overnight in LB (at the appropriate salinity) and then for either 3 or 17 h in fresh LB (or M9 medium at the same salinity, see below) for optimal growth or nutrient depletion[47], respectively (see Methods).

Exposing *E. coli* to nutrient depletion alone (i.e. 17 h growth in 10 g/L NaCl LB), favoured a steeper increase in intracellular 2-NBDG, compared to optimal growth conditions (i.e. 3 h growth in 10 g/L NaCl LB) with a mean fluorescence of 417 and 250 a.u. after 300 s incubation in 2-NBDG (Fig. 2b and 2a, respectively, ****, Table S2 and S3). However, nutrient depletion alone did not significantly affect 2-NBDG accumulation at steady state with a mean fluorescence of 740 and 733 a.u., respectively, at *t*=900 s (n.s., Table S2 and S3).

**Figure 2.**
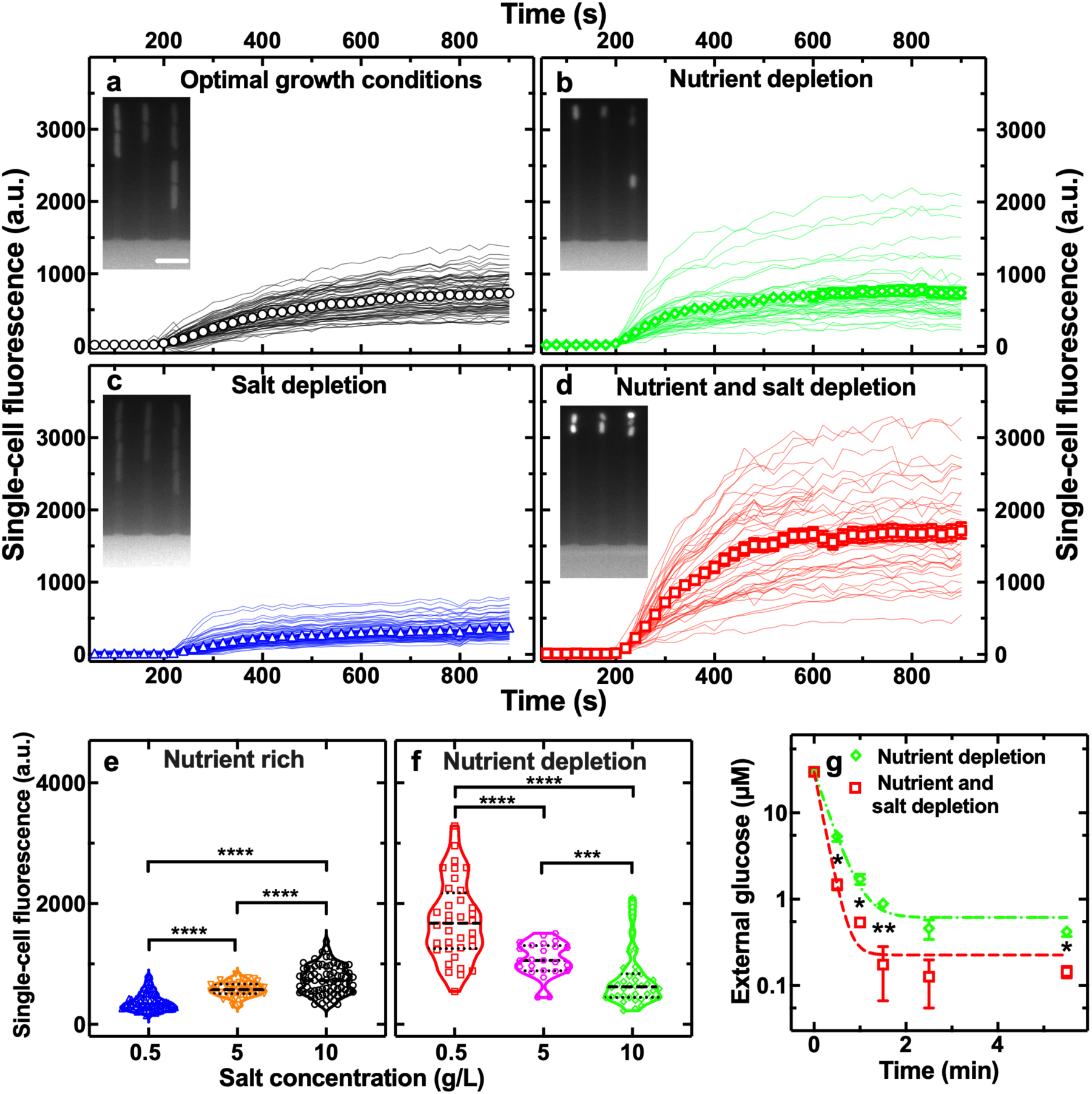
Glucose accumulation is greatest under combined nutritional and salinity depletion. Temporal dependence of the mean intracellular fluorescence of the glucose analogue 2-NBDG in individual *E. coli* under **a)** optimal growth conditions, **b)** nutrient depletion, **c)** salt depletion or **d)** combined nutrient and salt depletion. Lines are temporal dependences of the intracellular fluorescence of individual bacteria collated from biological triplicate. Symbols and error bars are the corresponding means and standard error of the means of such single-cell measurements. Noteworthy, we measured the 2-NBDG fluorescence as the mean fluorescent values of each pixel constituting each bacterium, thus normalizing by cell size. Coefficient of variations of these single-cell values and statistical tests of quantitative comparisons of values between different environments are reported in Table S2 and S3, respectively. Corresponding 2-NBDG intracellular fluorescence values during the removal of extracellular 2-NBDG are reported in Fig. S5. Insets: corresponding fluorescence images at *t*=900s when the intracellular 2-NBDG accumulation has reached saturation levels in individual bacteria. Measurements were carried out on N=76, 38, 90 and 46 individual bacteria, in a)-d), respectively. Salinity-dependent distribution of single-cell fluorescence after 900 s incubation in 2-NBDG in **e)** nutrient-rich or **f)** nutrient-depleted environments. Dashed and dotted lines indicate the median and quartiles of each distribution, respectively. **g)** Temporal dependence of the extracellular glucose concentration for *E. coli* under nutrient (green diamonds) or nutrient and salt depletion (red squares). N=3 biological replicates for each environment. The dashed dotted and dashed lines are one phase exponential decay fittings to the data yielding a significantly smaller time constant *Tau* for *E. coli* under combined nutrient and salt depletion compared to nutrient depletion alone (*Tau* = (9±1) s vs (17±1) s, respectively, *). *: p-value<0.05, **: p-value<0.01, ***: p-value<0.001, ****: p-value<0.0001.

Exposing *E. coli* to salinity depletion alone (i.e. 3 h growth in 0.5 g/L NaCl LB), caused a less steep intracellular 2-NBDG increase, compared to optimal growth conditions, with a mean fluorescence of 148 and 250 a.u. after 300 s incubation in 2-NBDG (Fig. 2c and 2a, respectively, ****, Table S2 and S3). Moreover, salt depletion also significantly reduced 2-NBDG accumulation at steady state with a mean fluorescence of 373 and 733 a.u., respectively, at *t*=900 s (****).

In striking contrast with the findings above, simultaneous exposure to nutritional and salinity depletion (i.e. 17 h growth in 0.5 g/L NaCl LB) caused a steeper intracellular 2-NBDG increase, compared to optimal growth conditions, with a mean fluorescence of 720 and 250 a.u. after 300 s incubation in 2-NBDG (Fig. 2d and 2a, respectively, ****, Table S2 and S3). Moreover, combined nutrient and salt depletion also significantly enhanced 2-NBDG accumulation at steady state with a mean fluorescence of 1714 and 733 a.u. at *t*=900 s, respectively (****, Table S2 and S3). These findings were further confirmed via separate flow cytometry measurements (red and black bars in Fig. S2, ****).

Moreover, we found that in nutrient-rich conditions 2-NBDG accumulation increased with salinity (Fig. 2e); on the other hand, under nutrient depletion 2-NBDG accumulation decreased with salinity (Fig. 2f). In fact, a 50% reduction in salinity (from 10 g/L down to 5 g/L NaCl) led to a 145% increase in 2-NBDG accumulation with a mean fluorescence of 740 and 1072 a.u. at *t*=900 s, respectively (***); a further 90% reduction in salinity (from 5 g/L down to 0.5 g/L NaCl, resembling the salinity change encountered during transition from the colon to the ileum[32]) led to a further 160% increase in 2-NBDG accumulation with a mean fluorescence of 1072 and 1714 a.u. at *t*=900 s, respectively (****). Taken together these data demonstrate that reducing the salinity content of the environment favours 2-NBDG accumulation in nutrient-poor but not in nutrient-rich environments.

Next, we verified that nutrient and salt depletion also synergistically boosts glucose accumulation (as a result of uptake and degradation), and not only the glucose analogue 2-NBDG. We performed plate reader based colorimetric assays on *E. coli* populations exposed to either nutrient, or simultaneous nutrient and salt depletion. We found that after 30 s incubation, the extracellular glucose concentration became significantly lower in *E. coli* that had been exposed to simultaneous nutrient and salt depletion compared to *E. coli* that had experienced nutrient depletion alone with a decay time constant *Tau* of (9.5±0.4) s and (16.7±1.2) s, respectively (*, red squares and green diamonds, respectively, in Fig. 2g). These data therefore confirm that *E. coli* accumulate glucose significantly faster and to higher levels after exposure to simultaneous nutrient and salt depletion compared to exposure to nutrient depletion alone.

We further confirmed that the findings above were not affected by molecular leakage through the cell membrane, cell integrity being essential for 2-NBDG uptake[5], by performing separate experiments using thioflavin T (ThT). This compound stains intracellular macromolecules[48, 49] and is of comparable size to 2-NBDG. We found that, in contrast to 2-NBDG, ThT accumulated to a significantly lesser extent in *E. coli* exposed to combined nutritional and salinity depletion compared to optimal growth conditions (Fig. S3b and S3a, red and black bars in Fig. S3c, respectively, and Table S4). Furthermore, *E. coli* exhibited similar growth curves in both LB formulations (i.e. 0.5 g/L or 10 g/L NaCl). This data confirmed that increased 2-NBDG accumulation under combined nutritional and salinity depletion was not due to molecular leakage through compromised bacterial membranes and that growth rate alone does not necessarily reflect important changes in bacterial metabolism[50]. Finally, we sought to rule out the possibility that increased glucose accumulation under combined nutritional and salinity depletion was a result of i) a nutritional shift from LB to M9 (used for growth and 2-NBDG measurements, respectively), ii) low abundance of divalent cations in LB medium or iii) changes in extracellular pH[45, 51]. In order to do so, we performed flow cytometry experiments on *E. coli* grown for 17 h in M9 with limited (i.e. 0.1 g/L) glucose or ammonium (i.e. carbon or nitrogen limitation, respectively,[52] see Methods) and either 0.5 or 10 g/L NaCl. Consistently with the data in Fig. 2, we found that *E. coli* exposed to combined nutritional and salinity depletion in M9 accumulated 2-NBDG to a significantly larger extent than *E. coli* exposed to nutritional depletion alone in M9 both with glucose (mean fluorescence of 3007 and 1045 a.u. at *t*=900 s, red and green violins, respectively, in Fig. S4a, ****) or ammonium as limiting factor (mean fluorescence of 3249 and 1776 a.u. at *t*=900 s, red and green violins, respectively, in Fig. S4b, ****). Furthermore, the measured extracellular pH values were the same (8.0±0.1) for the nutrient depleted and nutrient and salt depleted environments. Finally, *E. coli* displayed similarly large cell-to-cell differences in the accumulation of 2-NBDG after pre-culturing in LB or M9 (coefficient of variations of 38% and 60% after pre-culturing in 0.5 g/L or 10 g/L LB; coefficient of variations of 66% and 69% after pre-culturing in 0.5 g/L or 10 g/L glucose-limited M9; coefficient of variations of 43% and 64% after pre-culturing in 0.5 g/L or 10 g/L ammonia-limited M9). Therefore, the observed heterogeneity in 2-NBDG accumulation was not driven by the nutritional shift from LB to M9. Taken together this data demonstrate that when these two environmental changes come together, they manifest a synergistic effect on intracellular processes, such as glucose metabolism, that is greater than the effect of each environmental change alone.

### Glucose accumulation is heterogeneous under different environmental variations in a cell size-independent fashion

Besides the above described different 2-NBDG accumulation traits across variant environments, we also found substantial phenotypic heterogeneity within *E. coli* populations in the same environment. This heterogeneity increased in the presence of nutrient or salt depletion compared to optimal growth conditions (CV values of 2-NBDG intracellular fluorescence in Table S2), suggesting specialization in metabolic functions to endure environmental variations.

In order to gain a mechanistic understanding of these glucose accumulation traits in variant environments, we firstly investigated the role played by cell area in 2-NBDG accumulation. We studied the correlation of the total fluorescence of each bacterium at steady-state (*t*=900 s) with the area of each bacterium. We found strong positive correlation between single-cell area and total intracellular 2-NBDG fluorescence both via flow cytometry and via microfluidics-microscopy (Fig. 3a and 3b, respectively, Pearson correlation coefficient larger than 0.7 in all tested environments).

**Figure 3.**
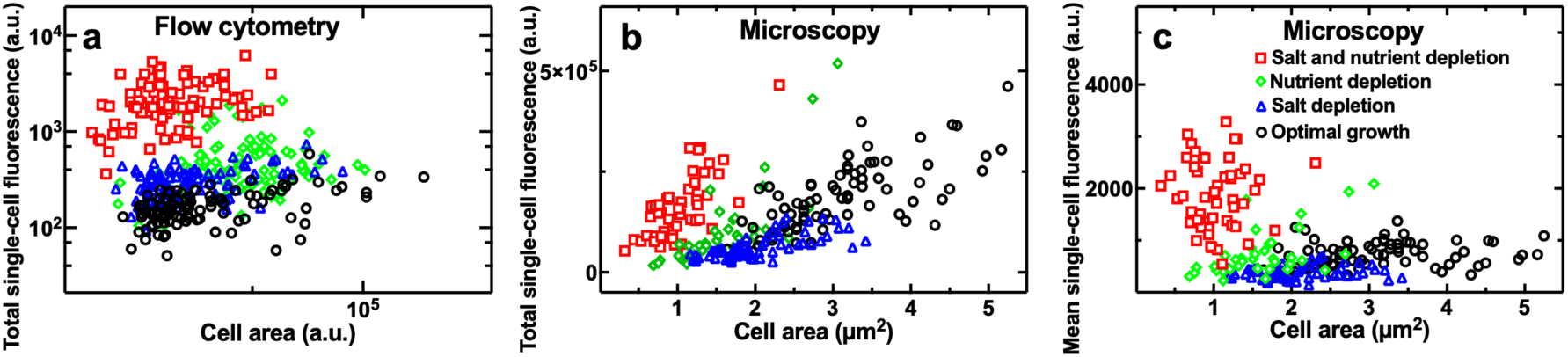
Glucose accumulation is heterogeneous across variant environments in a cell size-independent fashion. Size dependence of the total intracellular fluorescence of the glucose analogue 2-NBDG in individual *E. coli* under optimal growth conditions (circles), nutrient depletion (diamonds), salt depletion (triangles) or combined nutrient and salt depletion (squares) measured after 900s incubation in 2-NBDG via **a)** flow cytometry and **b)** microscopy in the mother machine. Pearson correlation coefficients of a) 0.35, 0.40, 0.42 and 0.32 and b) 0.71, 0.76, 0.69 and 0.71, respectively (all ****). **c)** Corresponding size dependence of the mean intracellular fluorescence normalized by cell size. Pearson correlation coefficients of 0.18 (ns), 0.57 (***), 0.16 (ns) and -0.01 (ns), respectively. ns=non significant, p-value>0.05; ***, p-value<0.001; ****, p-value<0.0001. N=100 representative measurements on individual bacteria in a) out of a total N=50000 measurements performed in biological triplicates. N>30 for each experimental condition collated from biological triplicates in b) and c).

However, when we calculated the mean 2-NBDG fluorescence for each bacterium, after normalizing by cell size[53], we found that, under combined nutritional and salinity depletion, *E. coli* exhibited significantly higher intracellular 2-NBDG fluorescence despite displaying significantly lower cell area compared to optimal growth conditions (squares and circles, respectively, in Fig. 3c). Moreover, we found a remarkable lack of correlation between the mean 2-NBDG fluorescence and cell size under optimal growth, salt depleted or nutrient and salt depleted conditions (Pearson correlation coefficients of 0.18 (ns), 0.16 (ns) and -0.01 (ns), respectively); we found instead a significantly positive correlation between 2-NBDG fluorescence and cell size under nutrient depletion (Pearson correlation coefficients of 0.57, ***). Finally, we still found substantial levels of heterogeneity in the mean 2-NBDG fluorescence values across the different environments (Fig. 3c). Taken together these data demonstrate that changes in cell size alone cannot account for the measured glucose accumulation traits and heterogeneities in variant environments in contrast to the current consensus[5, 15].

### Combined nutritional and salinity depletion boosts both glucose uptake and degradation

We then sought to decouple the contribution of uptake and degradation mechanisms on the measured 2-NBDG accumulation. To do this, we used a mathematical model describing 2- NBDG accumulation in single bacteria using ordinary differential equation 1 (see Methods). We inferred the uptake rate from the accumulation data in the presence of extracellular 2- NBDG (0<*t*<900s, Fig. 2) and obtained an independent estimate of the degradation rate from both the accumulation data in the presence of extracellular 2-NBDG (0<*t*<900s, Fig. 2) and the degradation data in the absence of extracellular 2-NBDG (*t*>1200s, Fig. S5).

Nutritional depletion caused a significant decrease in 2-NBDG uptake rate but also a significant increase in degradation rate compared to optimal growth conditions (Fig. 4a, 4d and 4e, Table S5). Moreover, both 2-NBDG uptake and degradation were more heterogeneous under nutritional depletion (Table S6), further suggesting specialisation in metabolic functions in stressed bacteria[44]. Finally, we did not find a significant correlation between uptake and degradation rates at the single-bacterium level under nutritional depletion (Pearson correlation coefficient of 0.27 compared to 0.42 for optimal growth conditions). This suggests that under nutritional depletion, some bacteria specialize in fast uptake at the expense of reduced degradation rates.

**Figure 4.**
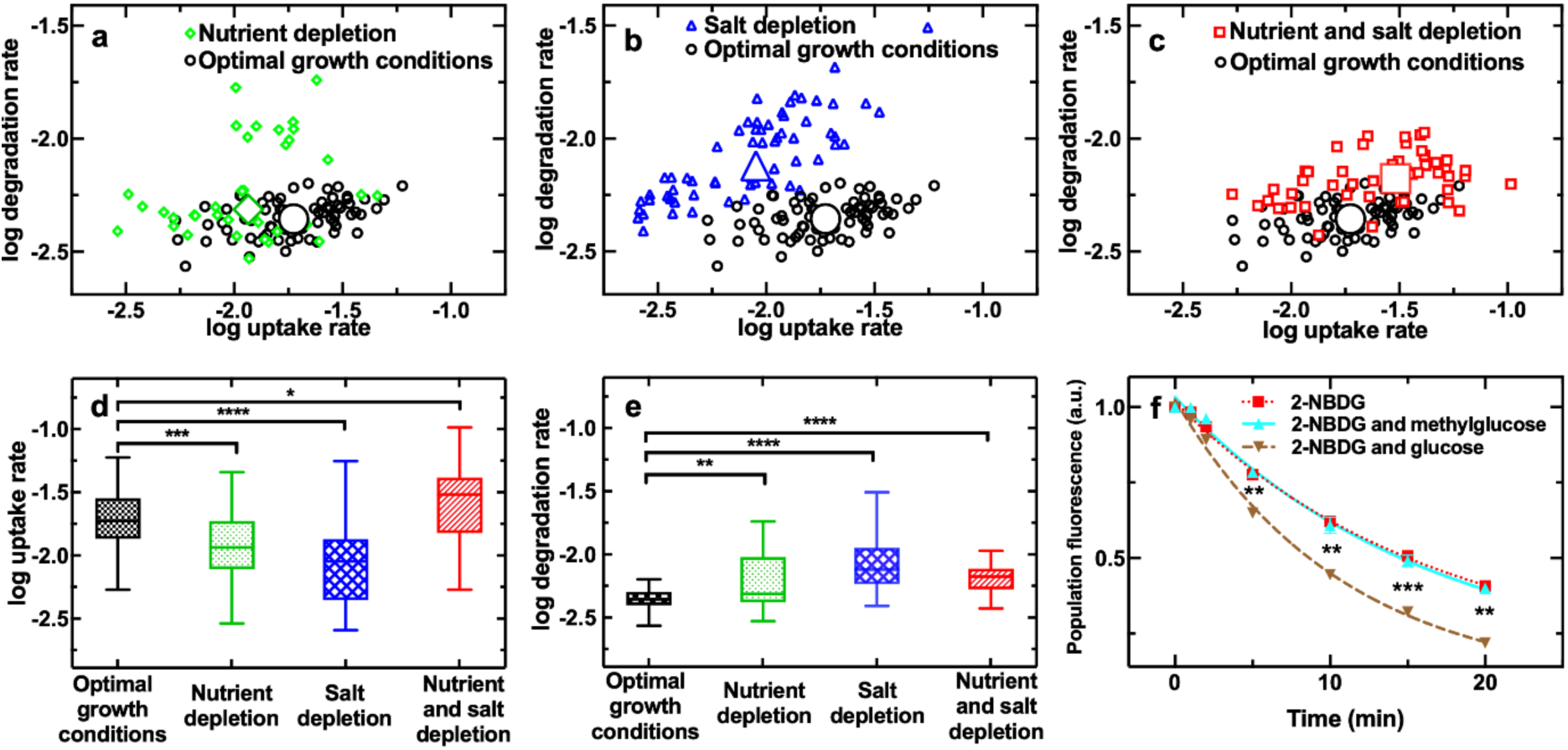
Combined nutritional and salinity depletion boosts both glucose uptake and degradation. **a)-c)** Single-cell correlation between the natural logarithm of 2-NBDG uptake rate and the natural logarithm of 2-NBDG degradation rate as predicted by Equation 1 for optimal growth conditions (circles), nutrient depletion (diamonds), salt depletion (triangles) or combined nutrient and salt depletion (squares), with Pearson correlation coefficients of 0.42 (***), 0.27 (ns), 0.80 (****) and 0.36 (*), respectively. N>30 for each experimental condition collated from biological triplicates, the large symbols are the medians of each set of single-cell values. Coefficient of variations of these single-cell values and statistical tests of quantitative comparisons of values between different environments are reported in Table S5 and S6, respectively. Corresponding distributions of **d)** natural logarithm of 2-NBDG uptake rate values and **e)** natural logarithm of 2-NBDG degradation rate values. The bottom and top of the box are the first and third quartiles, the band inside the box is the median, the bottom and top whiskers represent the 10th and 90th percentiles, respectively. **f)** Temporal dependence of the fluorescence of 2-NBDG when *E. coli* populations were incubated in 30 µM 2-NBDG alone (red squares), in the presence of 30 mM 3-O-methylglucose (cyan upward triangles), or in the presence of 30 mM glucose (brown downward triangles). N=3 biological replicates for each condition. The lines are one phase exponential decay fittings to the data yielding time constant *Tau* of (16.3 ± 1.1) min, (16.3 ± 1.2) min and (10.9 ± 0.5) min, respectively. *, p-value<0.05; ***, p-value<0.001; ****, p-value<0.0001.

Similarly, salinity depletion caused a significant decrease in 2-NBDG uptake rate but also a significant increase in degradation rate (Fig. 4b, 4d and 4e, Table S5). Furthermore, the CV of both uptake and degradation rate was higher under salinity depletion (Table S6). However, differently from nutritional depletion, salinity depletion favoured a strong positive correlation between uptake and degradation rates at the single bacterium level (Pearson correlation coefficient of 0.80). This suggests that salt depletion favours the emergence of a subset of the population specialising in both taking up and degrading 2-NBDG.

In contrast with the findings above, the combined presence of nutritional and salinity depletion significantly increased both the uptake and degradation rate as well as intra-population phenotypic heterogeneities in both parameters (Fig. 4c, 4d and 4e, Table S5 and S6). This environment yielded a significant correlation between uptake and degradation rate (Pearson correlation coefficient of 0.36). However, such correlation was weaker with respect to that measured in the presence of salt depletion alone. This suggests that the additional nutritional depletion favoured specialization in fast 2-NBDG uptake at the cost of reduced degradation rates compared to champion degraders exposed to salinity depletion alone.

Next, we set out to confirm that the degradation of 2-NBDG by *E. coli* is linked to activity of the bacterial glycolytic pathway, responsible for the degradation of glucose. We investigated the degradation of the fluorescence of 2-NBDG in the presence of a competitive and non-metabolizable inhibitor (i.e. 3-O-methyl glucose) or a competitive and metabolizable inhibitor (i.e. glucose). The presence of a metabolizable inhibitor would stimulate bacterial glycolytic activity and thereby enhance the 2-NBDG degradation, while the presence of a non-metabolizable inhibitor would have no effect[5]. Accordingly, we found that the degradation of the fluorescence of 2-NBDG in the presence of a competitive and metabolizable inhibitor was significantly faster than the degradation of the fluorescence of 2-NBDG in the absence of this inhibitor (brown downward triangles and red squares in Fig. 4f, respectively, time constant *Tau* of (10.9 ± 0.5) min and (16.3 ± 1.1) min, respectively) since such inhibitor promotes glycolytic activity. In contrast, we found that the degradation of the fluorescence of 2-NBDG in the presence of a competitive and non-metabolizable inhibitor was comparable to the degradation of the fluorescence of 2-NBDG in the absence of this inhibitor (cyan upward triangles and red squares in Fig. 4f, respectively, time constant *Tau* of (16.3 ± 1.2) min and (16.3 ± 1.1) min, respectively) since such inhibitor cannot be metabolized, thereby not promoting the bacterial glycolytic activity. These data therefore demonstrate that 2-NBDG is degraded as a consequence of the glycolytic pathway and can be used as a proxy for glucose degradation.

Taken together these data demonstrate that simultaneous nutritional and salinity depletion, that is the least favourable growth conditions investigated, has profound effects on glucose metabolism, increasing both uptake and degradation rates at the level of the individual bacterium, a synergistic effect that neither environmental changes alone can elicit.

### Molecular mechanisms underpinning glucose accumulation traits under nutrient or salt depletion

We then performed comparative transcriptomic and proteomic analysis between bulk *E. coli* cultures in the four different environments investigated and measured the log_2_ fold change in transcript or protein levels under nutrient or salt depletion compared to those measured in optimal growth conditions. We then restricted our combined transcriptomic and proteomic analysis to genes and proteins whose differential expression had a *p*-value adjusted for false discovery rate smaller than 0.05[54]. We found a significant correlation between gene and protein expression under salt depletion but no significant correlation under either nutrient or nutrient and salt depletion (Fig. 5a, Pearson coefficient R = 0.13, 0.06 and 0.07, **, ns, ns, respectively). We also found a stronger correlation between differential gene and protein regulation under nutrient and nutrient and salt depletion compared to salt and nutrient and salt depletion (Pearson coefficient R = 0.92 and 0.34, at the transcriptomic level and 0.89 and 0.45, at the proteomic level, all ****).

**Figure 5.**
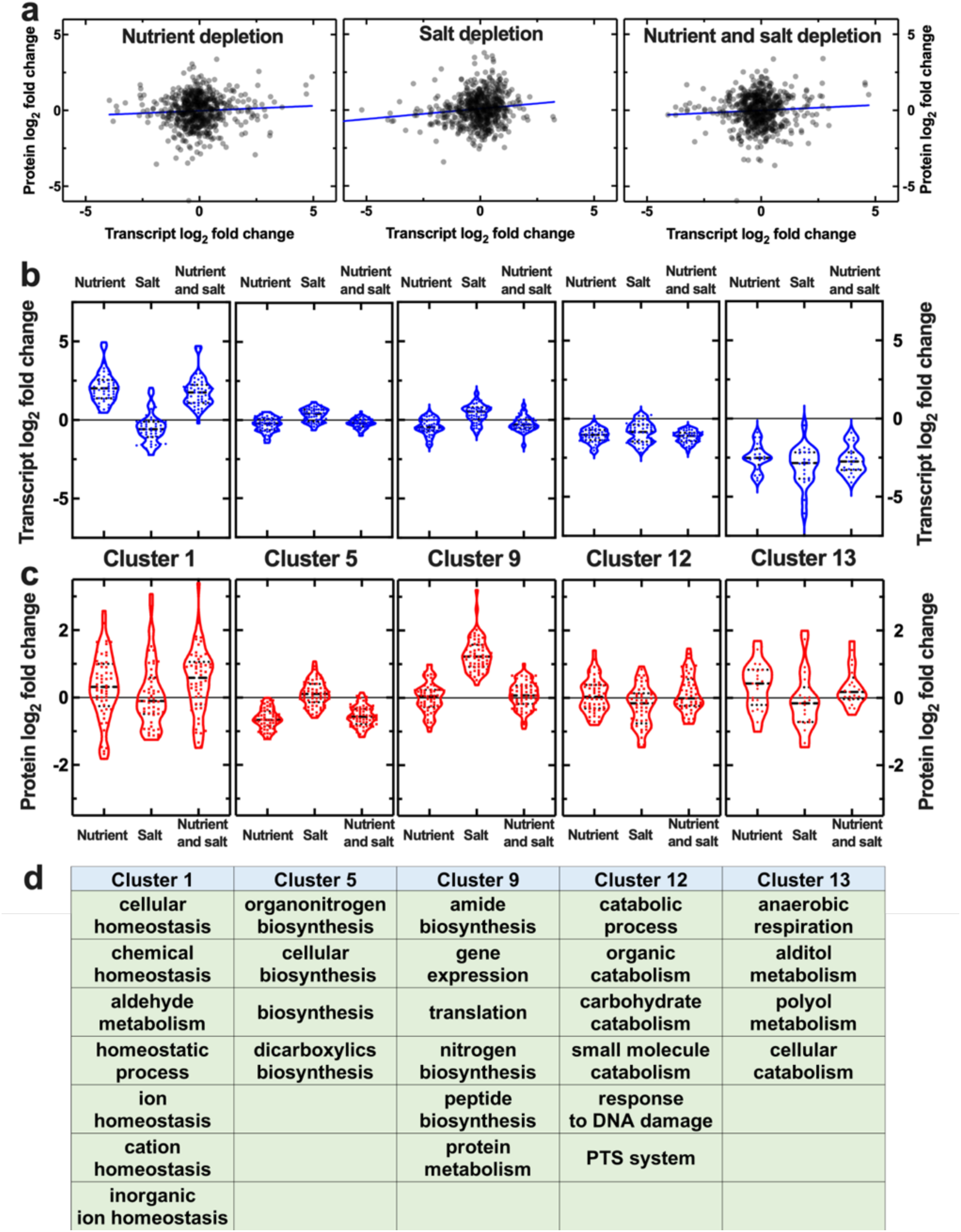
Molecular mechanisms underpinning glucose accumulation traits under nutrient or salt depletion. **a)** Correlation between the log_2_ fold change in gene and protein expression under nutrient, salt or nutrient and salt depletion. Each dot is the log_2_ fold change in the number of copies for a single gene or protein, blue lines are linear regressions to the data returning a Pearson correlation coefficient R=0.06, 0.13 and 0.07, (ns, ** and ns, respectively). Transcript and protein reads were measured via RNA-sequencing and proteomics on samples in biological triplicate and are reported in Supplementary files 2 and 3, respectively. Cluster analysis of the combined transcriptomic and proteomic data above returned 13 clusters with distinct patterns of **b)** gene and **c)** protein regulation under nutrient, salt or nutrient and salt depletion. Each dot represents the log_2_ fold change in the number of copies for a single gene or protein, dashed and dotted lines indicate the median and quartiles of each distribution, respectively, the grey solid lines indicate a log_2_ fold change of zero. The lists of genes belonging to each cluster are reported in Supplementary file 1. **d)** Biological processes significantly over-represented in each of the clusters above. For clarity, only clusters displaying significantly over-represented biological processes are reported in figure, data for the remaining clusters are reported in Supplementary file 1.

To determine biological processes underlying acclimation of sugar metabolism to changes in nutritional availability and environmental salinity, we clustered these combined transcriptomic and proteomic data, identifying 13 distinct patterns of gene and protein regulation (Fig. 5b and 5c, respectively, and Supplementary file 1).

Cellular biosynthetic processes (e.g. dicarboxylic acid, amide, organonitrogen and peptide biosynthesis), gene expression and translation were significantly downregulated at the transcriptomic and proteomic levels in both the nutrient depleted and the nutrient and salt depleted environments compared to salt depletion alone (Clusters 5 and 9 in Fig. 5b and 5c, ****).

Cell homeostasis biological processes including chemical, ion and cation homeostasis were significantly upregulated under both nutrient and nutrient and salt depletion compared to salt depletion alone or optimal growth conditions both at the transcriptomic and proteomic level (Cluster 1 in Fig. 5b and 5c, **). Anaerobic respiration and polyol metabolic processes were significantly upregulated under both nutrient and nutrient and salt depletion at the proteomic level compared to salt depletion alone or optimal growth conditions (Cluster 13 in Fig. 5c, *) in accordance with a previous report[55].

Crucially for this study, phosphoenolpyruvate-dependent sugar phosphotransferase systems (PTS) were significantly upregulated at the proteomic level under both nutrient and nutrient and salt depletion compared to salt depletion alone (Cluster 12 in Fig. 5c, **). These proteins included the PTS system galacticol- and mannose-specific EIID components GatA and ManZ, respectively, the latter transporting both mannose and glucose across the *E. coli* inner membrane[13]. Furthermore, glucose metabolic processes were upregulated at the proteomic level under nutrient or nutrient and salt depletion compared to salt depletion alone (Cluster 12 in Figure 5c, **). These proteins included the formate acetyltransferase 1 PflB that is involved in pyruvate fermentation (following glucose conversion to pyruvate) and formate synthesis; the L-lactate dehydrogenase LldD which converts pyruvate to lactate; the glucose-1-phosphatase Agp which converts glucose to glucose-1-phosphate; the putative glucose-6-phosphate epimerase YeaD; as well as catabolic processes of other sugars such as galacticol and deoxyribose.

To further validate these data, we performed clustering analysis on transcriptomic data alone and found that carbohydrate transport and metabolic processes were significantly downregulated under nutrient or nutrient and salt depletion compared to salt depletion alone or optimal growth conditions (Clusters 8 and 12 in Supplementary file 2, ****). In striking contrast, in our clustering analysis performed on proteomic data alone we found that carbohydrate metabolic processes were significantly upregulated under nutrient or nutrient and salt depletion compared to salt depletion alone or optimal growth conditions (Cluster 8 in Supplementary file 3, ****). Besides the proteins found in Cluster 12 of our combined transcriptomic-proteomic clustering analysis, Cluster 8 of the proteomic analysis included the PTS system glucose-specific EIIA component Crr; the phosphocarrier protein PtsH; the glucose-1-phosphate adenylyltransferase GlgC; the aconitate hydratase A AcnA; the enolase Eno; the mannose-6-phosphate isomerase ManA; the phosphoglycerate kinase Pgk; the pyruvate kinase PykF; the galactitol 1-phosphate 5-dehydrogenase GatD; the phosphoglucomutase Pgm that is a highly conserved enzyme which functions at a key point in glucose metabolism (the interconversion of glucose-1-phosphate and glucose-6-phosphate,[56]), it is involved in several processes including bacterial pathogenesis[57] and was previously found to be upregulated under nutrient depletion in minimal medium[55]. Here we found that Pgm was further upregulated under combined nutrient and salt depletion compared to nutrient or salt depletion alone (log_2_ fold change of 0.7, 0.4 and 0.3, respectively, Supplementary file 3) in accordance with our data on glucose degradation rates (Fig. 4e). Taken together these data demonstrate that under nutrient or combined nutrient and salt depletion glucose uptake and degradation are regulated at translation rather than transcription level.

Moreover, when we carried out phosphoproteomics on *E. coli* cultured in the four different environments above, we found that the phosphoglucomutase Pgm was one of the 13 proteins for which a phosphorylation event was detected (rows 8-9 in Supplementary file 4). Other phosphorylated proteins included PpsA and GpmM involved in gluconeogenesis and PpC and Icd (i.e. the first example of a protein phosphorylated on a serine or a threonine residue in bacteria[55]) involved in the tricarboxylic acid cycle. We measured phosphorylation at two different sites for the phosphopeptide GGPLADGIVITPSHNPPEDGGI that aligns with high confidence to the Pgm amino acid sequence. The first phosphorylation event was recorded on serine 13 of the phosphopeptide above with a site probability of 88% as defined by the phosphoRS node of Proteome Discoverer software (i.e. serine 146 in the full protein sequence in accordance with previous studies[55, 58]). The amino acid residues surrounding this serine are highly conserved in the phosphohexomutase superfamily to which Pgm belongs[59]. This stretch of residues is known as the phosphoserine loop and is involved in the transfer of a phosphoryl group from the serine residue above to glucose 6-phosphate to form a biphosphorylated sugar intermediate. A high level of phosphorylation of this serine residue increases Pgm flexibility and its enzymatic activity[59]. In accordance with our glucose degradation data (Fig. 4e), we measured higher Pgm phosphorylation levels at serine 13 under combined nutrient and salt depletion compared to optimal growth conditions (*, row 8 in Supplementary file 4). This indicates that the combined nutrient and salt depletion drives a 12- fold post-translational change as well as a 1.6-fold translational change in Pgm levels, thus explaining the higher glucose degradation rate reported in Fig. 4. Moreover, we measured significantly higher Pgm phosphorylation levels under combined nutrient and salt depletion compared to optimal growth conditions (*, row 9 in Supplementary file 4) at a second site within the phosphoserine loop. However, this site could not be defined with any degree of confidence (i.e. site probability < 70%). Finally, a significantly higher degree of phosphorylation under nutrient and salt depletion compared to optimal growth conditions was also measured for the phosphoenolpyruvate synthase PpsA and the isocitrate dehydrogenase Icd (* and **, rows 6 and 17 in Supplementary file 4) involved in gluconeogenesis and the tricarboxylic acid cycle, respectively.

Considering the significant variations in Pgm both at the translational and post-translational level, we set out to investigate the role played by Pgm in 2-NBDG accumulation. To do this, we performed single-cell 2-NBDG accumulation measurements on both a Δ*pgm* deletion mutant and the parental strain (PS). We found that under nutrient and salt depletion the Δ*pgm* deletion mutant displayed significantly lower 2-NBDG accumulation compared to the parental strain (purple hexagons and red squares in Fig. S6, respectively, ***). In contrast, under optimal growth conditions the Δ*pgm* deletion mutant displayed 2-NBDG accumulation comparable to the parental strain (magenta diamonds and black circles in Fig. S6, respectively, ns). These data further confirm that modifications of Pgm at the translational and post-translational level allow *E. coli* to enhance glucose accumulation.

Taken together these data demonstrate that simultaneous exposure to nutrient and salt depletion decreases gene expression, translation and biosynthetic processes, while increasing *E. coli* capability to take up and use glucose (and possibly other sugars) via variations at the translational and post-translational level but not at the transcriptional level, thus corroborating our glucose and 2-NBDG accumulation data presented in Fig. 2-4.

### Glucose accumulation is not regulated by molecules secreted in the environment

Finally, we also investigated the possibility that the combined nutritional and salinity depletion caused bacterial secretion of molecules that affect intracellular 2-NBDG accumulation. These molecules could include signalling secondary metabolites, such as putrescine and cadaverine, that affect the functioning of membrane transporters[54]. In order to test this hypothesis, we exposed *E. coli* grown in optimal growth conditions to the supernatant collected from *E. coli* exposed to combined nutritional and salinity depletion. After 1h exposure to such supernatant, we washed the microfluidic environment, introduced 30 µM 2-NBDG dissolved in glucose-free M9 medium and measured 2-NBDG accumulation in individual bacteria. We then compared these measurements to those performed without exposing bacteria to such supernatant (squares and circles, respectively, in Fig. S7). In contrast to the hypothesis above, we found that exposure to the supernatant collected from nutrient and salt depleted *E. coli* did not enhance 2-NBDG accumulation. Taken together these data suggest that the metabolic changes observed in the simultaneous nutritional and salinity depletion are not due to bacterial secretion of compounds that can alter glucose metabolism.

## DISCUSSION

Glucose uptake and utilization in bacteria have been previously linked to cell size, suggesting that both glucose uptake and intracellular conversion are maximal in favourable growth conditions, an idea that has led to the consensus that bacteria dwelling in stressful environments reduce their metabolic capabilities[15–17,60]. However, these findings were obtained either in optimal growth conditions or in the presence of only nutritional depletion[15]. In contrast, in natural environments such as soil, aquatic systems or the human body, microbial communities face multiple environmental changes[18, 35]. Therefore, it is crucial to predict the effects of multiple simultaneous environmental variations on the phenotypic diversity in microbial traits.

Here we demonstrate a hitherto unrecognised phenomenon in bacterial glucose metabolism by finding enhanced glucose accumulation traits when *E. coli* are simultaneously exposed to nutritional and salinity depletion. These traits are not displayed when bacteria are exposed to either depletion alone, suggesting that the effect of these changes on glucose metabolism is not additive[35]. In contrast with the general consensus, we show that these differences in glucose accumulation traits cannot be explained by differences in cell size or gene expression, a finding that needs to be taken into account when modelling metabolic fluxes and when designing bioproduction in *E. coli*[14] since we show that increasing cell size does not accelerate glucose metabolism.

We show instead that the measured metabolic changes are underpinned by variations in glucose transport and metabolism at the translational and post-translational level. Protein phosphorylation, especially on serine, threonine, or tyrosine, is one of the most common post-translational modifications in bacteria[55]. Protein phosphorylation controls cell metabolism and enhances cellular fitness under growth limiting conditions; for example, enzymes such as phosphoglycerate mutase, phosphoglucomutase and adenosine 5′-phosphosulfate kinase catalyse the turnover of phosphorylated sugars or metabolite phosphorylation by going through a phosphorylated intermediate state during catalysis[61]. A previous study found a global increase of protein phosphorylation levels under nutrient depletion[55]. Here we complement this understanding by demonstrating elevated protein phosphorylation levels under nutrient depletion, salt depletion or combined nutrient and salt depletion. Taken together these findings point to a likely role of protein phosphorylation in variant environments. Indeed, we found significantly higher phosphorylation levels of the phosphoglucomutase Pgm (at serine 146), a highly conserved enzyme[59], under nutrient and salt depletion compared to optimal growth conditions offering a mechanistic explanation of the measured glucose degradation rates. Furthermore, Pgm was upregulated at the translational level under nutrient depletion in accordance with a previous study using minimal medium[55], whereas we use LB medium, further confirming that this metabolic response is not dictated by the pre-culturing medium. We also found that Pgm was further upregulated under combined nutrient and salt depletion compared to nutrient depletion alone corroborating our data on glucose degradation rates. Finally, it is conceivable that other previously identified post-translational modifications (e.g. the acetylation of GapA and FbaA[58], both involved in carbohydrate degradation) could further contribute to the observed variations in glucose uptake and degradation rates.

We also showed that exposing *E. coli* growing in optimal conditions to the supernatant collected from cells under nutrient and salt depletion did not enhance glucose accumulation traits although this data should be corroborated in future via LC-MS metabolomics[62]. This finding demonstrates that enhanced glucose accumulation traits are not driven by the impact of physico-chemical properties of the nutrient and salt depleted environment on molecular transport, but rather by continuous cellular adaptation to such an environment.

These data corroborate previous work about the impact of salinity on carbon uptake in *Vibrio marinus* and cyanobacteria in natural environments[63, 64], and on the remodelling of *E. coli* glucose metabolism in the presence of environmental challenges[54, 65]. Furthermore, NaCl is routinely used in food products as an antimicrobial agent[66], therefore our findings that adding NaCl decreases glucose accumulation in stationary phase bacteria should be taken into account in both food preservation and bioproduction.

Our data also demonstrate, for the first time, that heterogeneity in glucose accumulation traits under combined nutrient and salt depletion cannot be ascribed to cell-to-cell differences in surface area alone neither can be ascribed to recovery from stationary phase[67]. In fact, we measured similar levels of heterogeneity in glucose uptake and degradation in exponentially growing and stationary phase *E. coli*. These data therefore suggest that cellular or environmental parameters other than cell size underpin heterogeneity in glucose metabolism adding to our current understanding of the relationship between cell growth rate and cellular processes[68–72], including those preparing a cell for surviving fluctuations in environmental conditions[73]. In this respect, we found that exposure to nutritional or salinity depletion increases heterogeneity in both glucose uptake and degradation. This could be explained by the recently reported heterogeneity in the expression of sugar metabolism genes[74, 75]. Taken together these data add new knowledge to the current understanding on phenotypic noise[76], corroborating the view that cell-to-cell differences are ubiquitous within microbial populations[67,76–78] and that there is substantial heterogeneity in the accumulation of metabolites[5,44,74,75,79–84] or antimicrobials[40,85,86].

This newly introduced experimental approach could be applied to other fields of research including medical mycology and crop protection, considering that glucose and its fluorescent analogue 2-NBDG is taken up by pathogenic fungi such as *Candida albicans*[87]. The newly discovered glucose accumulation traits should be considered when investigating cellular processes where salinity plays an important role such as cystic fibrosis associated lung infections[34] and the enteropathogens present in the ileum and colon[32] . If confirmed for microbes sampled from the environment, our findings will also inform modelling ecological and evolutionary dynamics considering the extensive impact of climate change on the salinity of freshwater ecosystems and naturally saline environments[25,30,35] which could consequently lead to a profound effect on the capabilities of some species to take up and use carbon sources.

## METHODS

### Strains, media and cell culture

All chemicals were purchased from Merck unless otherwise specified. Lysogeny Broth (LB, Melford) media made of 10 g/L Tryptone, 5 g/L Yeast extract and either 0.5, 5 or 10 g/L NaCl, were used for culturing *E. coli*. Noteworthy, the 5 and 10 g/L NaCl LB formulation are routinely used in microbiology, whereas the 0.5 g/L NaCl LB formulation is generally employed only for selective cultivation with antibiotics that require low salt conditions. The three formulations differ only in salt content, these do not differ in carbon and nitrogen source content that it is known to affect glucose metabolism[62]. LB agar plates of respective NaCl concentration with 15 g/L agar (Melford) were used for streak plates. Glucose-free M9-minimal media, used to wash cells and dilute 2-(N-(7-Nitrobenz-2-oxa-1,3-diazol-4-yl)Amino)-2-Deoxyglucose (2-NBDG) or Thioflavin T (ThT), was prepared using 5× M9 minimal salts (Merck), diluted as appropriate, with additional 2 mM MgSO_4_, 0.1 mM CaCl_2_, 3 µM thiamine HCl in milliQ water. Glucose or ammonia limited M9 media were prepared by adding to this solution 0.1 g/L glucose or 0.1 g/L NH_4_Cl, respectively[52], NaCl was added as required at a final concentration of 0.5 g/L or 10 g/L, as appropriate. The parental strain *E. coli* BW25113 and the single-gene knockout mutants Δ*pgm* were purchased from Dharmacon (GE Healthcare) and stored in 50% glycerol stock at -80°C. Streak plates for each strains were produced by thawing a small aliquot of the corresponding glycerol stock every 2 weeks and using LB agar containing either 0.5, 5 or 10 g/L NaCl. Exponentially growing cultures were obtained by inoculating 100 mL of LB (or M9 medium) of 0.5, 5 or 10 g/L NaCl content with 100 µl of stationary phase liquid culture *E. coli* BW25113, then placed in a shaking incubator at 200 rpm at 37°C for 3 hours. Overnight cultures were prepared inoculating a single colony of *E. coli* BW25113 in 200 mL of LB (or M9 medium) with 0.5, 5 or 10 g/L NaCl, then placed in a shaking incubator at 200 rpm at 37°C. Spent LB or M9 media used for resuspension of prepared bacteria in microfluidic assays was prepared by centrifugation of overnight cultures (4000 rpm, 20°C, 10 min). The supernatant was then double filtered (Medical Millex-GS Filter, 0.22 μm, Millipore Corp) to remove bacterial debris from the solution as previously reported[73]. 2-NBDG (Molecular weight=342 g/mol, ThermoFischer) was dissolved in dimethyl sulfoxide (DMSO) at a stock concentration of 10mg/ml and stored at -20°C. ThT (Molecular weight=319 g/mol, Merck) was dissolved in milliQ water at a stock concentration of 2 mM and stored at 4°C.

### Fabrication of microfluidic mother machine devices

Microfluidic mother machine devices were fabricated in polydimethylsiloxane (PDMS, Sylgard 184 silicone elastomer kit, Dow Corning) following previously reported protocols[73, 88]. A 10:1 (base:curing agent) PDMS mixture was cast on an epoxy mold of the mother machine device provided by S. Jun[41]. Each mother machine device contains approximately 6000 bacterial hosting channels with a width, height and length of 1, 1.5 and 25 µm, respectively. These channels are connected to a main microfluidic chamber with a width and height of 25 and 100 µm, respectively. After degassing the PDMS mixture, this was allowed to cure at 70°C for 2 hours, before unmoulding the solid PDMS replica from the mother machine epoxy mold. Using a 0.75 mm biopsy punch (RapidCore 0.75, Well-Tech) fluidic inlet and outlet accesses were created at the two ends of the main chamber of the mother machine. After ensuring accesses and surface of PDMS chip were completely clean using ethanol wash, N_2_ gas drying and removal of any small particles using adhesive tape (Scotch® Magic™ Tape, 3M), the PDMS chip, along with a glass coverslip (Borosilicate Glass No.1, Fisherbrand), were further cleaned and surfaces hydrophilized using air plasma treatment (10 s exposure to 30 W plasma power; Plasma etcher, Diener, Royal Oak, MI, USA) as previously reported[89]. Next, PDMS and glass were brought in contact and irreversibly bonded. Upon bonding, the device was placed for 5 min into an oven set at 70°C to enhance the adhesion between the PDMS and the glass surfaces. Finally, the device was filled with a 5 µL aliquot of 50 mg/ml bovine serum albumin (BSA) in milliQ water and incubated at 37°C for 1 hour; this step allowed to passivate the charge within the channels post oxygen plasma treatment, thus screening any electrostatic interaction between bacterial membranes and the glass and PDMS surfaces of the mother machine.

### Microfluidics-microscopy assay to measure 2-NBDG accumulation and degradation in individual E. coli bacteria

The bacterial inoculum to be injected in the mother machine device was prepared via centrifugation (5 min, 4000 rpm, 20°C) of a 50 ml aliquot of an exponentially growing or overnight culture (3 or 17 h after inoculation in LB or M9) and resuspended in spent LB (or M9) medium (both prepared as described above) to an OD_600_ of 75. 50 mg/mL BSA was added to the overnight culture before centrifugation to prevent bacterial cell aggregation[73]. The bacterial suspension was then injected into the mother machine device and incubated at 37°C aiming to fill each bacterial hosting channel at an average density of one cell. Subsequently, fluorinated ethylene propylene tubing (1/32” x 0.0008”) was inserted into both inlet and outlet accesses as previously reported[90]. A flow sensor unit (Flow Unit S, Fluigent, Paris, France) was connected to the inlet tubing. This flow sensor unit was also connected to a pressure control system (MFCS-4C, Fluigent), both being controlled via MAESFLO software (Fluigent) allowing for computerised, accurate regulation of fluid flow into the microfluidic device. The above described spent LB medium was used to clear the main channel of excess bacteria that had not entered the bacterial hosting channels, initially flowing at 300 μL/h for 8 min and then set to a flow rate of 100 μL/h. The loaded microfluidic device was mounted onto an inverted microscope (IX73 Olympus, Tokyo, Japan) following the flushing of the main channel with spent media and centred on one area of the mother machine containing 46 of the 6000 bacteria hosting channels; after visual inspection of the device, using automated stages for both coarse and fine movements (M-545.USC and P-545.3C7, Physik Instrumente, Karlsruhe, Germany), this area was selected on account of it containing the highest number of channels filled with one bacterium. Initial bright-field and fluorescence images were acquired using a 60×, 1.2 N.A. objective (UPLSAPO60XW, Olympus) and a sCMOS camera (Zyla 4.2, Andor, Belfast, UK), and, for the fluorescence image only, a FITC filter and exposure to blue LED illumination at 15% of its maximum power (200mW, CoolLED pE-300white, Andover, UK) for 0.1 s. Following this, 30 µM 2-NBDG in M9-minimal medium was introduced into the microfluidic device at an initial flow rate of 300 μL/h for 8 min, then 100 μL/h thereafter. Simultaneously, a fluorescence image was acquired every 20 s for a period of 900 s using the settings described above, a custom built LabView software and a 7-way Multi I/O timing cable (Andor, Belfast, UK) ensuring the LED only illuminated during image acquisition, thus reducing photobleaching. This same protocol was also employed to investigate the accumulation of 50 µM ThT in M9-minimal medium in individual *E. coli* bacteria.

At *t*=1200 s, the 2-NBDG solution was flushed away and exchanged to fresh LB or M9 medium (of appropriate NaCl concentration) flowing at an initial flow rate of 300 μl/h for 8 min, then 100 μl/h thereafter. This step allowed monitoring the degradation of 2-NBDG that had accumulated in each individual bacterium by acquiring a fluorescence image every 20 s until *t*=2100 s.

### Flow cytometry measurements

*E. coli* cultures were prepared as described above and 1 mL aliquoted into an Eppendorf at *t*=3 or 17 h and spun down at 13,400 rpm for 5 min using a microcentrifuge (SLS Lab Basics, UK). For uptake experiments, the bacterial pellet was resuspended in 1 mL of fluorescent substrate solution; either 30 µM 2-NBDG or 50 µM ThT and incubated for 15 or 45 min, respectively. After incubation, cells were pelleted at 13,400 rpm for 5 min using a table top centrifuge and resuspended in 1 ml of phosphate buffer solution (PBS) and further diluted as necessary. Measurements were then performed using a Beckman Coulter CytoFLEX S (Beckman Coulter, United States) and cell fluorescence quantified using the fluorescein isothiocyanate (FITC) channel (488 nm excitation and 525/40 nm band-pass filter), with PMT voltages of FSC 1000, SSC 500, FITC 250 and a threshold value of 10000 for SSC-A to exclude any background noise. These measurements were not normalized by cell size. Data were initially collected using CytExpert software, exported and later processed using GraphPad Prism 9. For degradation assays in the presence of competitive inhibitors, the bacterial pellet was resuspended in 30 µM 2-NBDG and incubated for 15 min as for the uptake experiments above. Following this 15-min incubation, 1 mL of D-glucose (metabolizable inhibitor) or 3-O-methylglucose (non-metabolizable inhibitor) was added to the sample at a final concentration of 30 mM. Samples were then diluted as necessary in PBS and measured via flow cytometry at intervals over 20 min as described above.

### Glucose colorimetric assay

17-hour stationary phase cultures grown in 0.5 g/L (nutrient and salt depletion) or 10 g/L NaCl LB (nutrient depletion) were prepared (as described above) in biological triplicate, reaching an optical density at 600 nm of 5 in both conditions. 1 mL of each culture was centrifuged at 13,400 rpm for 5 min using a microcentrifuge. Each pellet was subsequently resuspended in 1 mL 30 µM D-glucose and incubated for different time intervals. Following incubation, centrifugation was carried out as above but at 0°C, to reduce any further glucose uptake by cells. The supernatant of each sample was collected while the pellet was discarded. Using a glucose assay kit (Sigma Aldrich, Montana, United States), 50 µL of each supernatant sample was added to 50 µL master reaction mix (46 µL glucose assay buffer, 2 µL glucose probe and 2 µL glucose enzyme mix) and incubated in the dark at room temperature for 15 min. Oxidisation of any glucose present in the sample occurred during incubation, thus generating a colorimetric product, the absorbance of which was measured at 570 nm using a CLARIOstar PLUS plate reader (BMG Labtech, UK). For each experimental repeat, glucose standards were obtained by performing a serial dilution between 0 and 10 µL of a 1 nmole/µL glucose standard solution added to 50 µl of the master reaction mix above, brought to 100 µL per well with glucose assay buffer as needed. Absorbances collected for glucose standard wells were then used as a standard curve by which sample absorbances were compared and interpolated. To calculate the concentration of glucose present in each sample, background absorbance (assay blank of standard curve where 0 µl of glucose standard solution was present in the sample) was first subtracted and then the following equation was used:

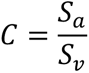

Where *S_a_* is the amount of glucose in the unknown sample (in nmole) from standard curve, *S_v_* is the sample volume (µL) added into the well and *C* describes the concentration of glucose in the sample.

### Image and data analysis

Images were processed using ImageJ software as previously described[88, 91] tracking each individual bacterium throughout their incubation in and removal of 2-NBDG. A rectangle was drawn around each bacterium at every time point, obtaining its width, length and mean fluorescence intensity. The mean fluorescence intensity for each bacterium was normalised by cell size, to account for cell cycle related variations in glucose accumulation[53]. The background fluorescence (i.e. the fluorescence of extracellular 2-NBDG) at each time point was measured for each bacterium as follow as follows: a rectangle, of similar dimensions to those drawn around the bacteria, was drawn and positioned, at the same distance from the main channel, in the nearest channel that did not contain bacteria and the mean fluorescence value within this rectangle was extracted via ImageJ. This background fluorescence was subtracted from the corresponding bacterium’s fluorescence value at every time point as previously reported[73].

These fluorescence data were then analysed and plotted using GraphPad Prism 9. Statistical significance was tested using unpaired, two-tailed, Welch’s *t*-test unless specified otherwise. Error bars displayed in all graphs represent the standard error of the mean (SEM), however, due to the large sample sizes, error bars in some figures are small compared to the corresponding mean values and thus hidden behind data points. Pearson correlation, medians and coefficient of variations were also calculated using GraphPad Prism 9.

### Transcriptomic analysis

RNA isolation, library preparation, sequencing, and transcriptomic data processing was performed as previously reported[54, 92]. Briefly, *E. coli* cultures in high or low salt exponential (optimal growth conditions or salt depletion, respectively) and stationary phase (nutrient depletion or nutrient and salt depletion, respectively) were prepared as described above and 500 µL aliquots were taken from each culture after *t*=3 and 17 h after inoculation in LB, respectively, in biological triplicate for each of the four environmental conditions. RNAprotect Bacteria Reagent (Qiagen) was added to each aliquot. Extractions were performed using RNeasy Mini Kit (Qiagen) for exponential phase aliquots and RNeasy Micro Kit (Qiagen) for stationary phase aliquots following manufacturer instructions and previously reported protocols[54, 92]. DNA removal during extraction was carried out by using RNase-Free DNase I (Qiagen). RNA concentration and quality were measured using Qubit 1.0 fluorometer (ThermoFisher Scientific) and 2200 TapeStation (Agilent), respectively, and only samples with an RNA integrity number larger than 8 were taken forward. Transcript abundance was quantified using Salmon for each gene in all samples. Subsequent differential analysis was performed using DEseq2 in R software to quantify the log_2_ fold change in transcript reads[93] for each gene in aliquots under nutritional, salinity or combined nutritional and salinity depletion each compared to growth in optimal conditions. Significantly differentially expressed genes were defined as having a log_2_ fold change with respect to optimal growth conditions greater than 1 and a *p*-value adjusted for false discovery rate of <0.05[24].

### Proteomic Analysis

*E. coli* whole cell preparation was performed using 1.5 L cultures as described above. At 3 and 17 h, for exponential and stationary phase respectively, each culture was centrifuged at 4700 rpm ThermoFisher, USA) at 4°C for 20 min. Pellets were then resuspended in 5 mL of lysis buffer (20 mM Tris-HCl (pH 8.0), 0.5 M NaCl). Subsequently, one cOmplete™, EDTA-free Protease Inhibitor Cocktail (Roche, Sigma Aldrich) tablet and one phosSTOP™ (Roche, Sigma Aldrich) tablet were added to each sample. Sonication cycles for complete cell lysis were performed at 70% amplitude in a Sonics Vibracell VC-505 instrument (Sonics and Materials Inc., Newton, CT, USA) on ice. Lysate was then centrifuged to remove cell debris at 14400 rpm 4°C for 30 min and pellet was discarded. Aliquots of 80 µg of each sample were digested with trypsin (2.5 µg trypsin per 100 µg protein; 37°C, overnight), labelled with Tandem Mass Tag (TMT) eleven plex reagents according to the manufacturer’s protocol (Thermo Fisher Scientific) and the labelled samples pooled.

For the Total proteome analysis, an aliquot of 50 µg of the pooled sample was desalted using a SepPak cartridge according to the manufacturer’s instructions (Waters, Milford, Massachusetts, USA). Eluate from the SepPak cartridge was evaporated to dryness and resuspended in buffer A (20 mM ammonium hydroxide, pH 10) prior to fractionation by high pH reversed-phase chromatography using an Ultimate 3000 liquid chromatography system (Thermo Fisher Scientific). In brief, the sample was loaded onto an XBridge BEH C18 Column (130 Å, 3.5 µm, 2.1 mm × 150 mm, Waters, UK) in buffer A and peptides eluted with an increasing gradient of buffer B (20 mM Ammonium Hydroxide in acetonitrile, pH 10) from 0-95% over 60 min. The resulting fractions (5 in total) were evaporated to dryness and resuspended in 1% formic acid prior to analysis by nano-liquid chromatography tandem mass spectrometry (LC MSMS) using an Orbitrap Fusion Lumos mass spectrometer (Thermo Scientific).

For the Phospho proteome analysis, the remainder of the TMT-labelled pooled sample was also desalted using a SepPak cartridge (Waters, Milford, Massachusetts, USA). Eluate from the SepPak cartridge was evaporated to dryness and subjected to TiO_2_-based phosphopeptide enrichment according to the manufacturer’s instructions (Pierce). The flow-through and washes from the TiO_2_-based enrichment were then subjected to FeNTA-based phosphopeptide enrichment according to the manufacturer’s instructions (Pierce). The phospho-enriched samples were again evaporated to dryness and then resuspended in 1% formic acid prior to analysis by nano-LC MSMS using an Orbitrap Fusion Lumos mass spectrometer (Thermo Scientific).

High pH RP fractions (Total proteome analysis) or the phospho-enriched fractions (Phospho-proteome analysis) were further fractionated using an Ultimate 3000 nano-LC system in line with an Orbitrap Fusion Lumos mass spectrometer (Thermo Scientific). In brief, peptides in 1% (vol/vol) formic acid were injected onto an Acclaim PepMap C18 nano-trap column (Thermo Scientific). After washing with 0.5% (vol/vol) acetonitrile 0.1% (vol/vol) formic acid, peptides were resolved on a 250 mm × 75 μm Acclaim PepMap C18 reverse phase analytical column (Thermo Scientific) over a 150 min organic gradient, using 7 gradient segments (1-6% solvent B over 1 min, 6-15% B over 58 min, 15-32%B over 58 min, 32-40%B over 5 min, 40-90%B over 1 min, held at 90%B for 6 min and then reduced to 1%B over 1 min) with a flow rate of 300 nL/min. Solvent A was 0.1% formic acid and Solvent B was aqueous 80% acetonitrile in 0.1% formic acid. Peptides were ionized by nano-electrospray ionization at 2.0 kV using a stainless-steel emitter with an internal diameter of 30 μm (Thermo Scientific) and a capillary temperature of 300°C.

All spectra were acquired using an Orbitrap Fusion Lumos mass spectrometer controlled by Xcalibur 3.0 software (Thermo Scientific) and operated in data-dependent acquisition mode using an SPS-MS3 workflow. Fourier-transformed mass-spectrometry 1 (FTMS1) spectra were collected at a resolution of 120,000, with an automatic gain control (AGC) target of 200,000 and a maximum injection time of 50 ms. Precursors were filtered with an intensity threshold of 5,000 according to charge state (to include charge states 2-7) and with monoisotopic peak determination set to Peptide. Previously interrogated precursors were excluded using a dynamic window (60 s +/-10 ppm). The MS2 precursors were isolated with a quadrupole isolation window of 0.7m/z. Ion trap mass-spectrometry (ITMS2) spectra were collected with an AGC target of 10 000, max injection time of 70ms and Collision induced dissociation (CID) energy of 35%.

For FTMS3 analysis, the Orbitrap was operated at 50,000 resolution with an AGC target of 50,000 and a maximum injection time of 105 ms. Precursors were fragmented by high energy collision dissociation (HCD) at a normalised collision energy of 60% to ensure maximal TMT reporter ion yield. Synchronous Precursor Selection (SPS) was enabled to include up to 10 MS2 fragment ions in the FTMS3 scan.

The raw data files were processed and quantified using Proteome Discoverer software v2.1 (Thermo Scientific) and searched against the UniProt *Escherichia coli* (strain K12) [83333] database (downloaded November 2020: 8061 entries) using the SEQUEST HT algorithm. Peptide precursor mass tolerance was set at 10 ppm, and MS/MS tolerance was set at 0.6 Da. Search criteria included oxidation of methionine (+15.9949) as a variable modification and carbamidomethylation of cysteine (+57.0214) and the addition of the TMT mass tag (+229.163) to peptide N-termini and lysine as fixed modifications. For the Phospho-proteome analysis, phosphorylation of serine, threonine, tyrosine, histidine and aspartic acid (+79.966) was also included as a variable modification. Searches were performed with full tryptic digestion and a maximum of 2 missed cleavages were allowed. The reverse database search option was enabled and all data was filtered to satisfy false discovery rate (FDR) of 5%. Protein differential abundance analysis was carried out using *limma*[94], which has been shown to be appropriate for proteomics data[95]. *p*-values were adjusted for false discovery rate using the method of Benjamini and Hochberg[96].

### Cluster and gene ontology analysis

Clustering analysis was performed using the mclust package (version 5.4.7) for R[97]. Clustering was performed on the transcriptomic and proteomic log_2_ fold-change data separately and then simultaneously on both data sets. Hyperspherical models of equal and unequal volume were tested for a range of 2 to 20 clusters and the minimal, best-fitting model was identified by the Bayes information criterion. We observed generally larger fold-change magnitudes in the transcriptomic data, which may dominate in simultaneous clustering on both data sets. Therefore, prior to simultaneous clustering of proteomic and transcriptomic data, the fold-changes for each condition within each ‘omics data set were standardized by dividing by the sample standard deviations.

Gene Ontology enrichment analysis was performed using the clusterProfiler package (version 3.16.1) for R[98]. Enrichment in terms belonging to the “Biological Process” ontology was calculated for each gene cluster, relative to the set of all genes quantified in the experiment, via a one-sided Fisher exact test (hypergeometric test). *P* values were adjusted for false discovery by using the method of Benjamini and Hochberg[96]. Finally, the lists of significantly enriched terms were simplified to remove redundant terms, as assessed via their semantic similarity to other enriched terms, using clusterProfiler’s simplify function.

### Mathematical model and parameter inference

In order to rationalise the data acquired via our microfluidics-microscopy assay, we described the dynamics of intracellular 2-NBDG concentration in single cells using the following ordinary differential equation (ODE):

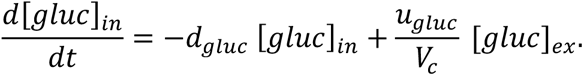

This equation describes the change of intracellular 2-NBDG concentration, [*gluc*]*_in_*, over time (left hand-side term) as the sum of two processes: intracellular 2-NBDG degradation (negative term in the right hand-side) occurring at rate *d_gluc_* [*gluc*]*_ in_* , and 2-NBDG uptake (positive term in the right hand-side) that we model as a first-order kinetics occurring at rate 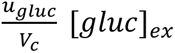, similarly to previous models^5^. Furthermore, [*gluc*]*_ex_* denotes the time-varying, extracellular 2-NBDG concentration. Here, since we measure the fluorescent intensity in the bacterial hosting channels (dashed line in Fig. 1) we chose to solve the ODE numerically approximating [*gluc*]*_ex_* at all times as a piecewise linear function. We note that with any functional form of [*gluc*]*_ex_* the ODE admits to an analytical solution, however the complex structure of the solution (involving time integration) defeats the purpose of using this directly for inference. An alternative approach would have been to assume instantaneous changes of external 2-NBDG at times *t* = 0 (addition) and *t* = *t*, = 1,200*s* (removal) and use the following simple analytical solution for inference purposes:

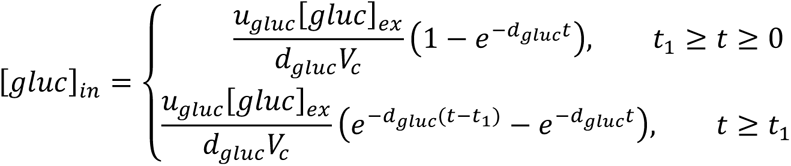

However, such an approach misses out information (i.e. non-instantaneous 2-NBDG delivery) that is crucial for providing an accurate description of the kinetics in 2-NBDG accumulation in single cells. Rate constant *u_gluc_*(*μ*m^2^s^-1^) and *d_gluc_*(s^-1^) dictate the rate of 2-NBDG uptake, and degradation, respectively. Parameter *V_c_* (*μ*m^2^) denotes the cellular volume, and its presence in the denominator of the uptake rate term ensures that intracellular 2-NBDG concertation is reduced in cells with larger volumes due to dilution (other parameters being the same). We estimate *V_c_* using the measured width and length of each cell and assuming a rod-like shape. We use the model and Bayesian inference techniques to extract information regarding rate parameters *u_gluc_*, *d_gluc_*from our single-cell measurements of intracellular 2-NBDG accumulation and degradation. To capture cell-to-cell heterogeneity we employ a hierarchical Bayes approach where parameters *u_gluc_*and *d_gluc_*vary between single cells according to an underlying population distribution. To model this unknown population distribution we use an infinite Gaussian mixture model (iGMM)[99], which being a non-parametric Bayesian model offers greater modelling flexibility than standard parametric distributional models. We note that the hierarchical Bayes approach also allow us to naturally incorporate in the model uncertainty (or lack of information) regarding the prior distribution of *u_gluc_* and *d_gluc_*, as we do not have to specify explicit priors for these parameters but instead specify priors for the hyper-parameters of the iGMM model. Choices of priors for the iGMM hyper-parameters are given in Table 1. To sample from the posterior distribution of the two parameters, *u_gluc,i_* and *d_gluc,i_*, for each cell *i*, we employed an iterative Gibbs sampling scheme. Each iteration of the sampling scheme, indexed by *k*, consists of two steps. In the first step, parameters 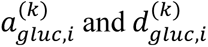 are sampled for each cell (indexed by *i*) given the data, ***y****_i_* = {*y_i,t_*: *i* = 1, … , *M*, *t* = 1, … , *Z*}, cellular volume *V_c,i_* , external 2-NBDG profile [*gluc*]*_ex,i_*, coefficient of variation of the measurement error *CV_error_* , and the iGMM parameterisation from the previous iteration, 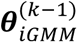. In the second step, iGMM parameters, 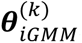, are sampled anew given the current values of 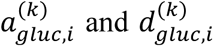. In summary, the algorithm involves sampling iteratively from the following target distributions:

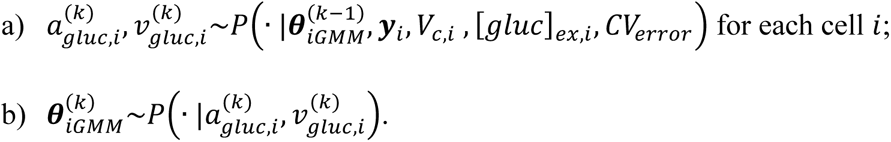

To sample parameters *u_gluc,i_* and *d_gluc,i_* in step a) we assume that the 2-NBDG accumulation and degradation measurement, *y_i,t_*, taken form cell *i* at time-point *t* obeys a gaussian distribution, 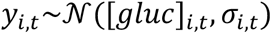, with mean [*gluc*]*_i,t_* and standard deviation 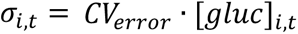. Here [*gluc*]*_i,t_* is obtained by solving the model numerically using the corresponding rate parameters, cellular volume, and external 2-NBDG concertation profile. This assumption allows us to use a single step of the Metropolis-Hastings algorithm to draw samples from the target distribution since this is proportional to a product of densities (gaussians and mixture of gaussians) all of which can be straightforwardly evaluated, i.e.,

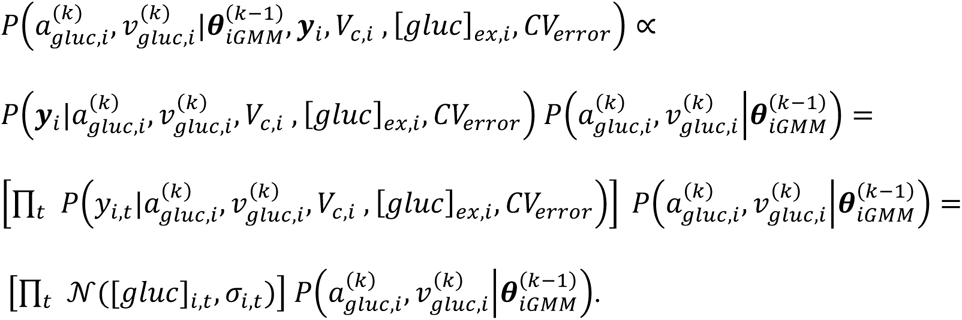

In step b) of the algorithm, we sampled from the target distribution using the algorithm as proposed by Rasmussen[99]. In the first iteration of the scheme, *k* = 1, we initialise 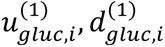 to values obtained using maximum likelihood estimation (perfomed using Matlab’s in-built nonlinear least-squares solver lsqcurvefit; using the Levenberg-Marquardt optimization algorithm and with the maximum number of iterations set to 15). The scheme was implemented and run using Matlab R2018.

**Table 1.** Hyperparameters of the infinite Gaussian mixture model and the associated priors used in our analysis. *μ*_ϑ_ , Σ_ϑ_ are obtained in the first step of our algorithm as the mean and covariance matrix of the model parameters values, 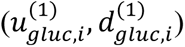, obtained via maximum likelihood estimation.

| Hyper-parameters | Hyper parameter description | Prior |
| --- | --- | --- |
| $\alpha$ | Concentration parameter of the Dirichlet prior. | $\alpha \sim \text{Gamma}(1,1)$ |
| $\lambda$ | Mean of the 2D-Guasssian prior for component means | $\lambda \sim \text{Normal}(\mu_{\vartheta}, \Sigma_{\vartheta})$ |
| $r^{-1}$ | St. deviation of the 2D-Guasssian prior<br>for component means. | $r \sim \text{Wishart}(\Sigma_{\theta}^{-1}/2, 2)$ |
| $2\beta$ | Shape parameter of the 2D Wishart<br>prior for component precisions (ie,<br>inverse st. devs.) | $\beta - 1 \sim \text{Gamma}(1, 1)$ |
| $w^{-1}/2$ | Scale matrix of the 2D Wishart prior<br>for component precisions (ie, inverse<br>st. devs.) | $w \sim \text{Wishart}(\Sigma_{\theta}/2, 2)$ |

## ACKNOWLEDGMENTS

G.G. was supported by an EPSRC DTP PhD studentship (EP/M506527/1). M.V. and K.T.A. gratefully acknowledge financial support from the EPSRC (EP/N014391/1). U.L. was supported through a BBSRC grant (BB/V008021/1) and an MRC Proximity to Discovery EXCITEME2 grant (MCPC17189). This work was further supported by a Royal Society Research Grant (RG180007) awarded to S.P. and a QUEX Initiator grant awarded to S.P. and K.T.A.. D.S.M., T.A.R. and S.P.’s work in this area is also supported by a Marie Skłodowska-Curie project SINGEK (H2020-MSCA-ITN-2015-675752) and the Gordon and Betty Moore Foundation Marine Microbiology Initiative (GBMF5514). B.M.I. acknowledges support from a Wellcome Trust Institutional Strategic Support Award to the University of Exeter (204909/Z/16/Z). This project utilised equipment funded by the Wellcome Trust Institutional Strategic Support Fund (WT097835MF), Wellcome Trust Multi User Equipment Award (WT101650MA) and BBSRC LOLA award (BB/K003240/1).

## AUTHOR CONTRIBUTIONS

S.P. designed the research and developed the project. G.G. and U.L. performed the experiments. M.V. and K.T.A. developed and implemented the mathematical model. B.M.I. performed the clustering and gene ontology analysis. D.S., P.O. and K.M performed the transcriptomics analysis. S.R. and D.S.M. assisted G.G. during protein extraction, flow cytometry and colorimetric assays. K.H. performed the proteomics and phosphoproteomics analysis. G.G., M.V., U.L., B.M.I., D.S., N.N., D.S.M., S.R., K.H., P.G.P. T.A.R. and S.P. analysed the data. G.G. and S.P. wrote the paper. All authors read and approved the final manuscript.

## SUPPLEMENTAL INFORMATION

**Figure S1.**
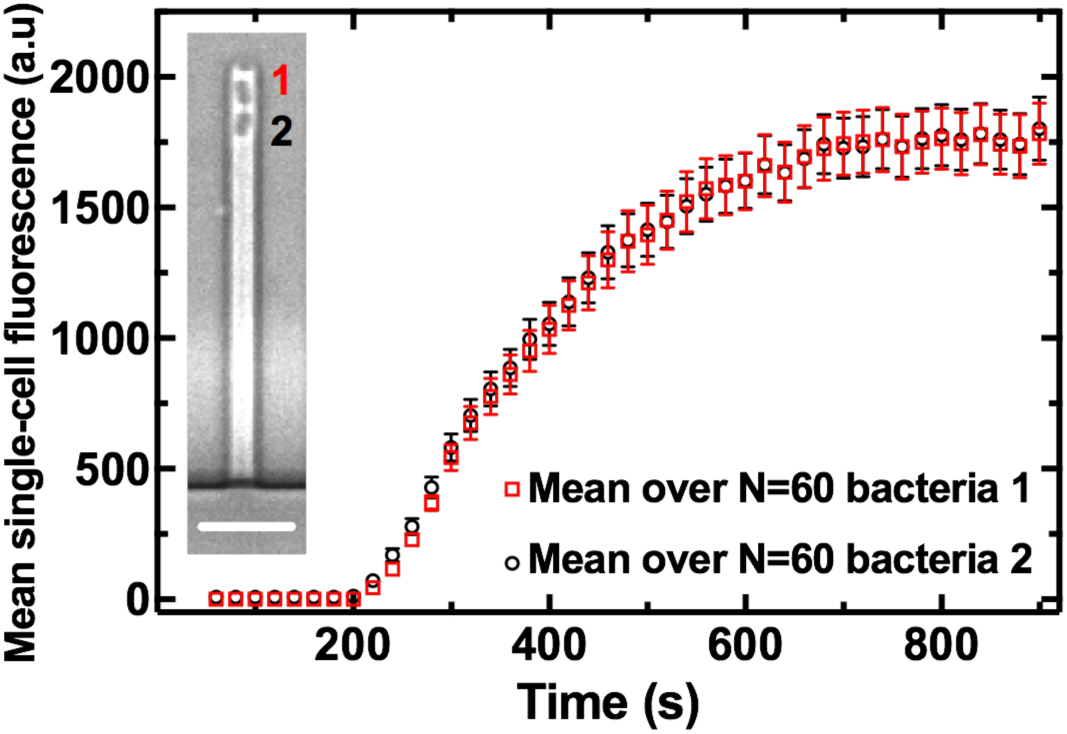
Temporal dependence of the mean intracellular fluorescence of the glucose analogue 2-NBDG averaged over 60 individual *E. coli* at the top of each dead-end bacteria-hosting channel (red squares, position 1) or below such cells (black circles, position 2). Bacteria at position 2 are closer to the main microfluidic chamber i.e. the 2-NBDG source, whereas bacteria at position 1 are screened by one cell. Statistical comparisons are reported in Table S1. Inset: bright-field microscopy image illustrating bacteria in positions 1 and 2, scale bar: 5 µm.

**Table S1.** Statistical comparisons of 2-NBDG accumulation in E. coli at the top of each dead-end bacteria-hosting channel or below such cells (positions 1 and 2 in Fig. S1, respectively).

| Time (s) | t ratio | df | p-value |
| --- | --- | --- | --- |
| 0 |  |  |  |
| 100 | 1,26 | 58 | 0,21 |
| 200 | 1,60 | 58 | 0,11 |
| 300 | 0,52 | 58 | 0,60 |
| 400 | 0,18 | 58 | 0,86 |
| 500 | 0,12 | 58 | 0,91 |
| 600 | 0,01 | 58 | 0,99 |
| 700 | 0,11 | 58 | 0,92 |
| 800 | 0,07 | 58 | 0,94 |
| 900 | 0,11 | 58 | 0,91 |

**Table S2.** Temporal dependence of the mean and coefficient of variation of intracellular fluorescence of the glucose analogue 2-NBDG over at least 30 individual *E. coli* (collated from biological triplicate) per environmental condition as detailed in Figure 2.

| Time (s) | Optimal growth conditions |  | Nutrient depletion |  |
| --- | --- | --- | --- | --- |
|  | Mean fluorescence (a.u.) | CV fluorescence (%) | Mean fluorescence (a.u.) | CV fluorescence (%) |
| 0 | 0 | 0 | 0 | 0 |
| 100 | 17 | 48 | 22 | 61 |
| 200 | 37 | 90 | 40 | 49 |
| 300 | 250 | 46 | 417 | 58 |
| 400 | 435 | 35 | 546 | 58 |
| 500 | 533 | 33 | 647 | 55 |
| 600 | 613 | 32 | 692 | 57 |
| 700 | 685 | 32 | 762 | 55 |
| 800 | 699 | 33 | 769 | 58 |
| 900 | 733 | 32 | 740 | 60 |
| 1000 | 730 | 34 | 683 | 54 |
| 1100 | 761 | 32 | 663 | 59 |
| 1200 | 580 | 46 | 458 | 54 |
| 1300 | 350 | 47 | 293 | 61 |
| 1400 | 231 | 49 | 208 | 64 |
| 1500 | 156 | 52 | 150 | 67 |
| 1600 | 112 | 52 | 113 | 69 |
| 1700 | 83 | 55 | 91 | 67 |
| 1800 | 63 | 55 | 76 | 69 |
| 1900 | 53 | 53 | 64 | 64 |
| 2000 | 42 | 47 | 51 | 64 |
| Time (s) | Salt depletion |  | Nutrient and salt depletion |  |
|  | Mean fluorescence (a.u.) | CV fluorescence (%) | Mean fluorescence (a.u.) | CV fluorescence (%) |
| 0 | 0 | 0 | 0 | 0 |
| 100 | 12 | 86 | 12 | 86 |
| 200 | 13 | 84 | 16 | 90 |
| 300 | 148 | 70 | 722 | 43 |
| 400 | 262 | 45 | 1208 | 42 |
| 500 | 298 | 43 | 1507 | 40 |
| 600 | 332 | 39 | 1669 | 38 |
| 700 | 326 | 41 | 1648 | 39 |
| 800 | 317 | 43 | 1676 | 38 |
| 900 | 373 | 39 | 1714 | 38 |
| 1000 | 393 | 43 | 1583 | 39 |
| 1100 | 429 | 38 | 1670 | 38 |
| 1200 | 328 | 45 | 1146 | 46 |
| 1300 | 155 | 38 | 582 | 55 |
| 1400 | 83 | 50 | 341 | 64 |
| 1500 | 44 | 98 | 186 | 55 |
| 1600 | 34 | 118 | 124 | 60 |
| 1700 | 23 | 128 | 82 | 79 |
| 1800 | 25 | 143 | 53 | 70 |
| 1900 | 29 | 151 | 40 | 62 |
| 2000 | 18 | 169 | 27 | 71 |

**Table S3.**
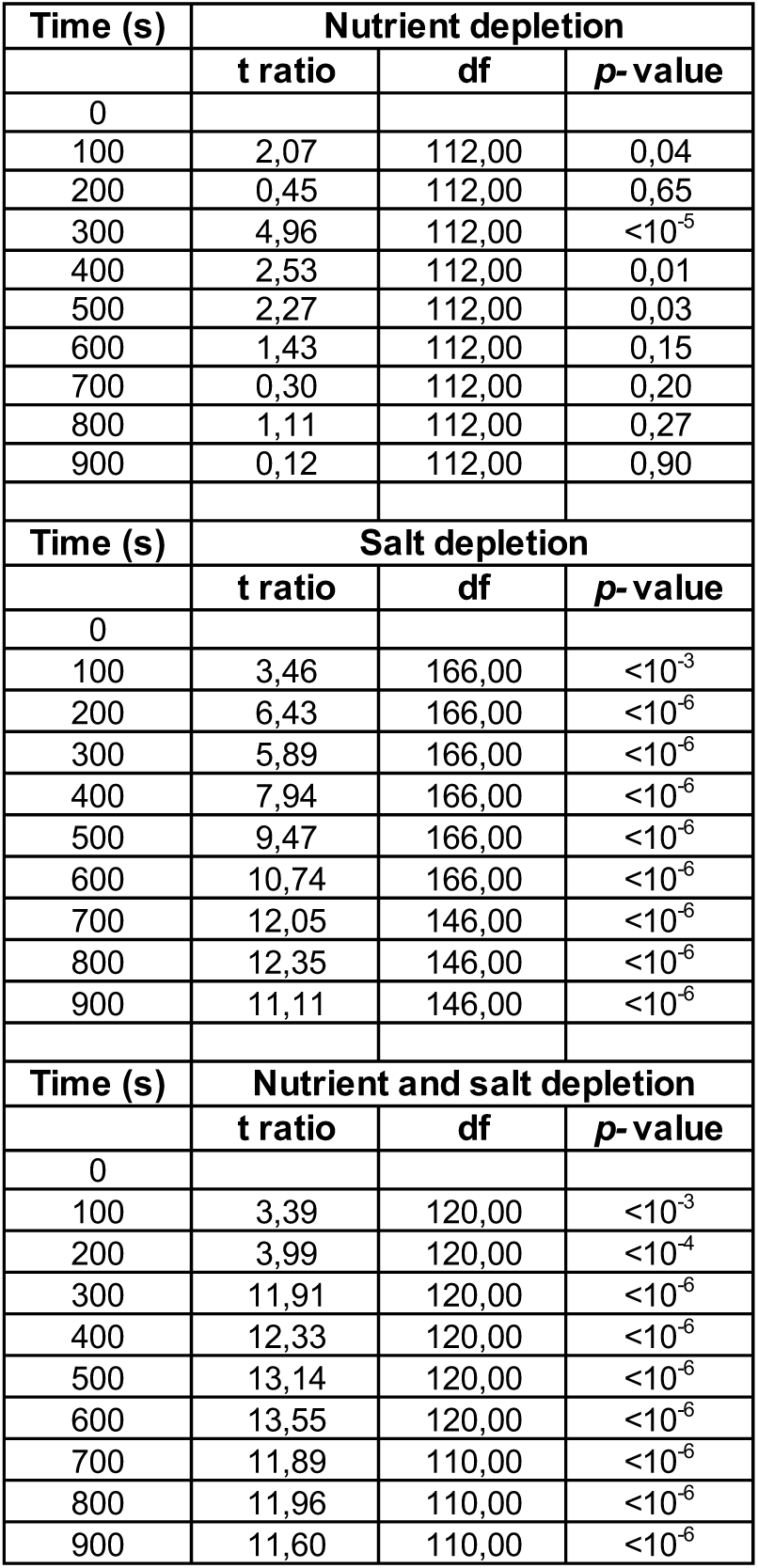
Statistical comparisons of 2-NBDG accumulation in E. coli under nutrient, salt or combined nutrient and salt depletion compared to optimal growth conditions.

| Time (s) | Nutrient depletion |  |  |
| --- | --- | --- | --- |
|  | t ratio | df | p-value |
| 0 |  |  |  |
| 100 | 2,07 | 112,00 | 0,04 |
| 200 | 0,45 | 112,00 | 0,65 |
| 300 | 4,96 | 112,00 | <10<sup>-5</sup> |
| 400 | 2,53 | 112,00 | 0,01 |
| 500 | 2,27 | 112,00 | 0,03 |
| 600 | 1,43 | 112,00 | 0,15 |
| 700 | 0,30 | 112,00 | 0,20 |
| 800 | 1,11 | 112,00 | 0,27 |
| 900 | 0,12 | 112,00 | 0,90 |
| Time (s) | Salt depletion |  |  |
|  | t ratio | df | p-value |
| 0 |  |  |  |
| 100 | 3,46 | 166,00 | <10<sup>-3</sup> |
| 200 | 6,43 | 166,00 | <10<sup>-6</sup> |
| 300 | 5,89 | 166,00 | <10<sup>-6</sup> |
| 400 | 7,94 | 166,00 | <10<sup>-6</sup> |
| 500 | 9,47 | 166,00 | <10<sup>-6</sup> |
| 600 | 10,74 | 166,00 | <10<sup>-6</sup> |
| 700 | 12,05 | 146,00 | <10<sup>-6</sup> |
| 800 | 12,35 | 146,00 | <10<sup>-6</sup> |
| 900 | 11,11 | 146,00 | <10<sup>-6</sup> |
| Time (s) | Nutrient and salt depletion |  |  |
|  | t ratio | df | p-value |
| 0 |  |  |  |
| 100 | 3,39 | 120,00 | <10<sup>-3</sup> |
| 200 | 3,99 | 120,00 | <10<sup>-4</sup> |
| 300 | 11,91 | 120,00 | <10<sup>-6</sup> |
| 400 | 12,33 | 120,00 | <10<sup>-6</sup> |
| 500 | 13,14 | 120,00 | <10<sup>-6</sup> |
| 600 | 13,55 | 120,00 | <10<sup>-6</sup> |
| 700 | 11,89 | 110,00 | <10<sup>-6</sup> |
| 800 | 11,96 | 110,00 | <10<sup>-6</sup> |
| 900 | 11,60 | 110,00 | <10<sup>-6</sup> |

**Figure S2.**
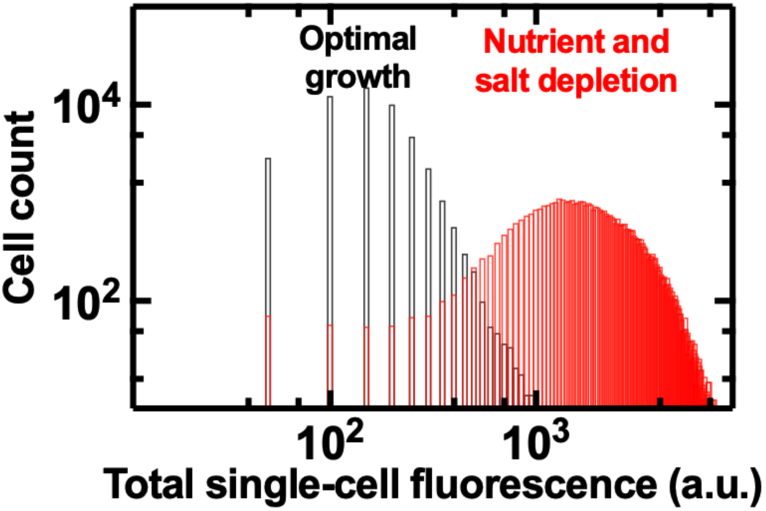
Total intracellular fluorescence of the glucose analogue 2-NBDG in individual *E. coli* under optimal growth conditions and combined nutrient and salt depletion (black and red bars, respectively) measured by flow cytometry after 900s bulk incubation in 2-NBDG. Noteworthy, these measurements were not normalized by cell size.

**Figure S3.**
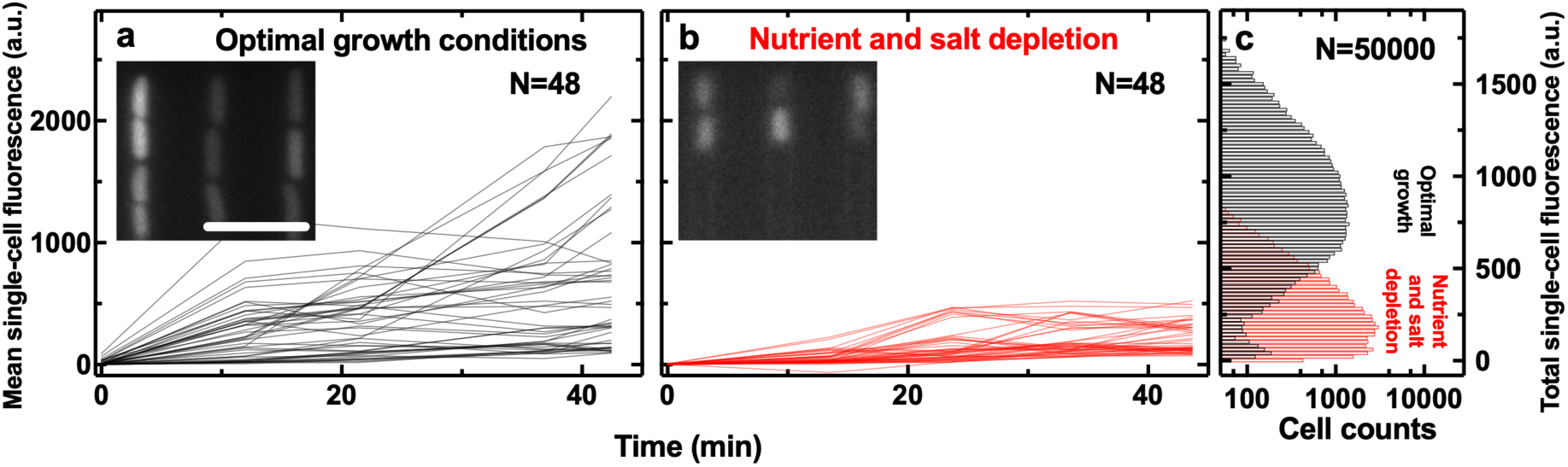
Temporal dependence of the mean intracellular fluorescence of thioflavin T in individual *E. coli* in **a)** optimal growth conditions and **b)** under combined nutrient and salt depletion. Lines are temporal dependences of the intracellular fluorescence of individual bacteria collated from biological triplicate. Noteworthy, we measured thioflavin T fluorescence as the mean fluorescent values of each pixel constituting each bacterium, thus normalizing by cell size. Insets: corresponding fluorescence images at t=45min when the intracellular ThT accumulation has reached saturation levels in individual bacteria. **c)** Corresponding total intracellular fluorescence of thioflavin T under optimal growth conditions or combined nutrient and salt depletion (black and red bars, respectively) measured by flow cytometry after 45min bulk incubation in thioflavin T. Noteworthy, these measurements were not normalized by cell size. Statistical comparisons are reported in Table S4.

**Table S4.** Statistical comparisons of thioflavin T accumulation in E. coli in optimal growth conditions and under combined nutrient and salt depletion as measured via single-cell microfluidics-microscopy and flow cytometry.

| Time (min) | Microfluidics-microscopy assay |  |  | Flow cytometry assay |  |  |
| --- | --- | --- | --- | --- | --- | --- |
|  | t ratio | df | p-value | t ratio | df | p-value |
| 0 | 3,78 | 94 | $<10^{-6}$ | | | |
| 12 | 7,98 | 94 | $<10^{-6}$ | | | |
| 22 | 5,56 | 94 | $<10^{-6}$ | | | |
| 37 | 6,01 | 94 | $<10^{-6}$ | | | |
| 42 | 6,23 | 94 | $<10^{-6}$ | 257,9 | 99578 | $<10^{-6}$ |

**Figure S4.**
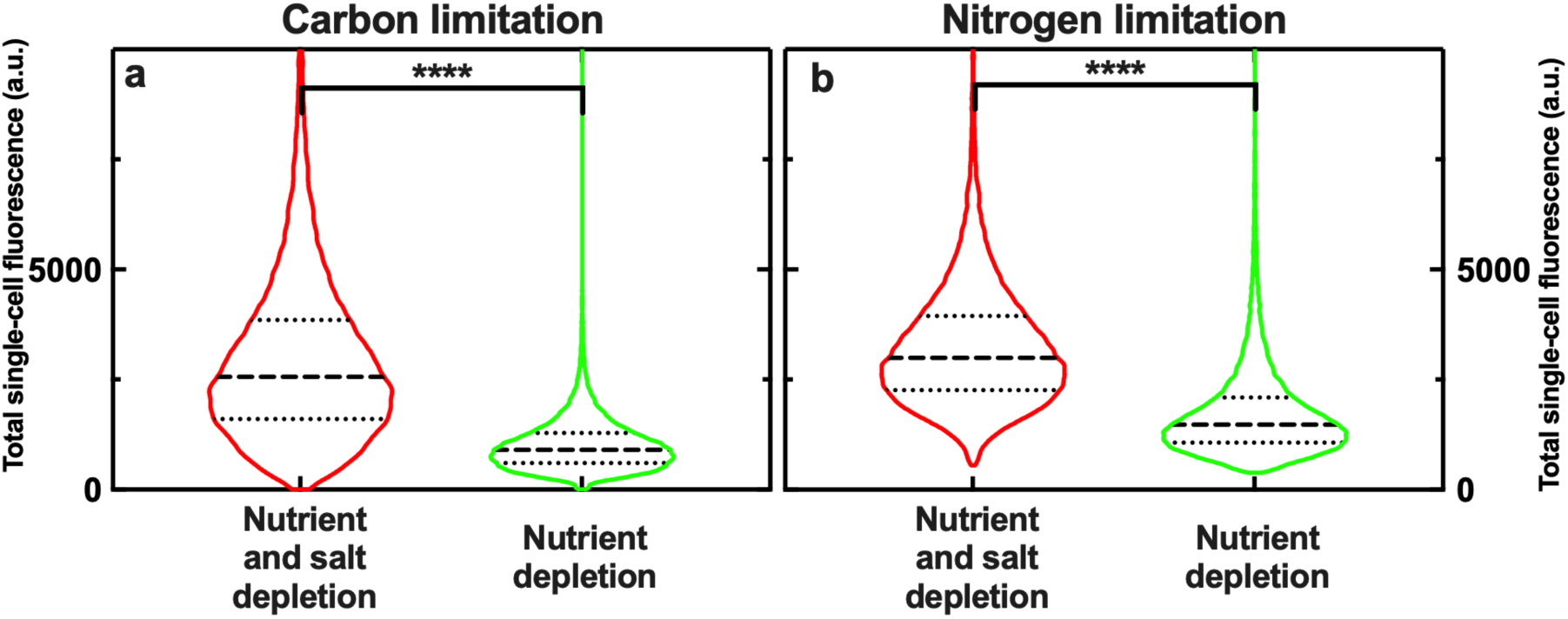
Distribution of total intracellular fluorescence of 2-NBDG under nutrient depletion alone or combined nutrient and salt depletion (green and red violins, respectively) using M9 minimal medium with limited (i.e. 0.1 g/L) **a)** glucose or **b)** ammonia. Measurements were performed on 50,000 bacteria for each environmental condition using flow cytometry after 900s bulk incubation in 2-NBDG. These measurements were not normalized by cell size. ****: p-value <0.0001.

**Figure S5.**
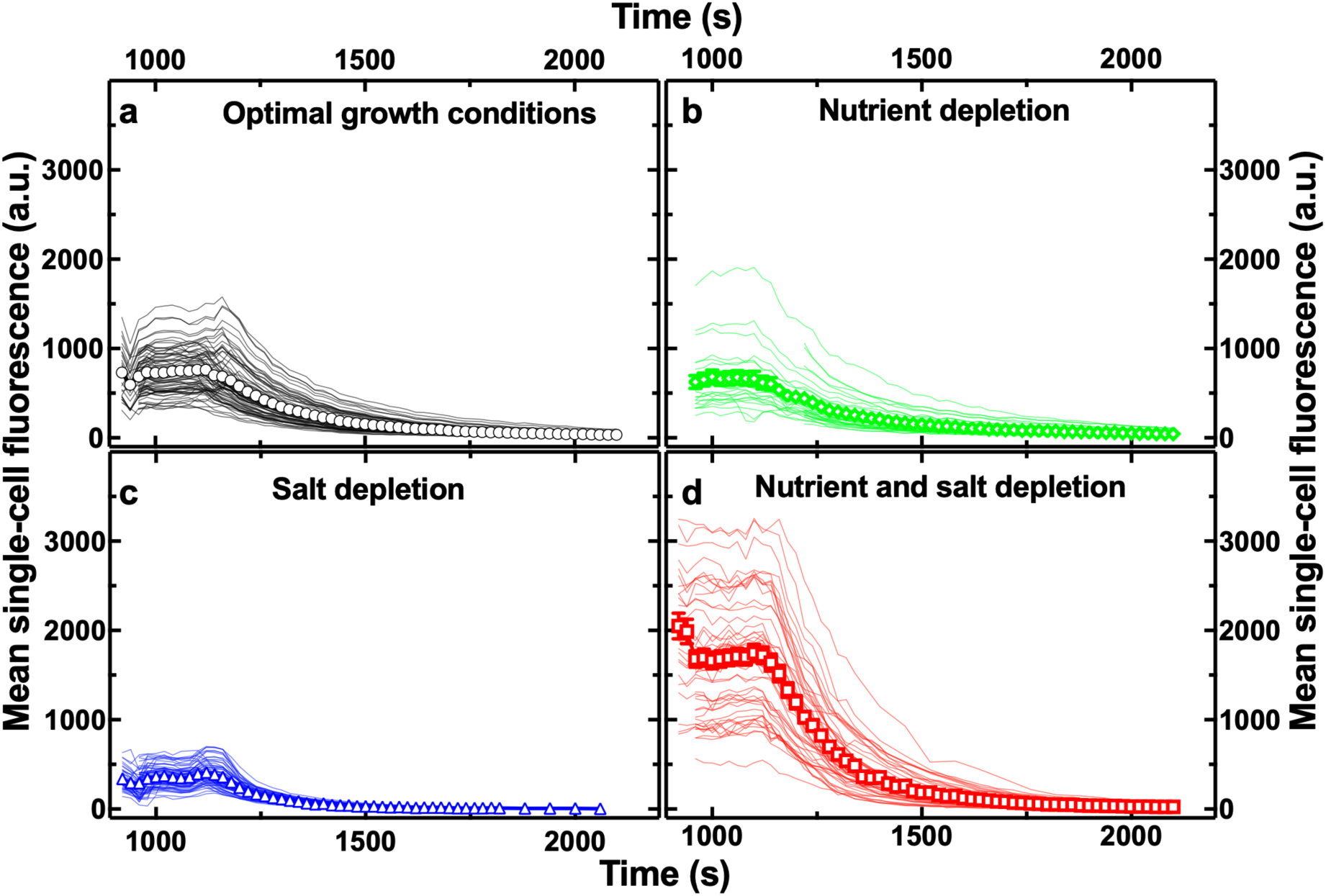
Temporal dependence of the mean intracellular fluorescence of the glucose analogue 2-NBDG in individual *E. coli* under **a)** optimal growth conditions, **b)** nutrient depletion, **c)** salt depletion or **d)** combined nutrient and salt depletion during removal of 2-NBDG from the extracellular environment. Lines are temporal dependences of the intracellular fluorescence of individual bacteria from biological triplicate. Symbols and error bars are the corresponding means and standard error of the means of such single-cell measurements. Means and coefficient of variations of these single-cell values are reported in Table S2. These measurements were normalized by cell size. Measurements were carried out on N=76, 38, 90 and 46 individual bacteria, in a)-d), respectively.

**Table S5.** Statistical comparisons of the predicted 2-NBDG uptake and degradation values under nutritional, salinity or combined nutritional and salinity depletion compared to optimal growth conditions.

| Comparisons with optimal growth | Uptake rate |  |  | Degradation rate |  |  |
| --- | --- | --- | --- | --- | --- | --- |
|  | t ratio | df | p-value | t ratio | df | p-value |
| Nutrient depletion | 3.88 | 62,0 | 0.0003 | 3.41 | 42 | 0.0015 |
| Salt depletion | 7.46 | 116,0 | <0.0001 | 10.97 | 83 | <0.0001 |
| Nutrient and salt depletion | 2.59 | 86,0 | 0.01 | 9.30 | 80 | <0.0001 |

**Table S6.** Median and coefficient of variation (CV) of the predicted uptake and degradation rate values in optimal growth conditions, under nutritional, salinity or combined nutritional and salinity depletion.

| Environment | Uptake rate |  | Degradation rate |  |
| --- | --- | --- | --- | --- |
|  | Median | CV (%) | Median | CV (%) |
| Optimal growth conditions | -1,73 | 13 | -2,35 | 3 |
| Nutrient depletion | -1,94 | 15 | -2,31 | 9 |
| Salt depletion | -2,05 | 15 | -2,12 | 9 |
| Nutrient and salt depletion | -1,52 | 19 | -2,18 | 5 |

**Figure S6.**
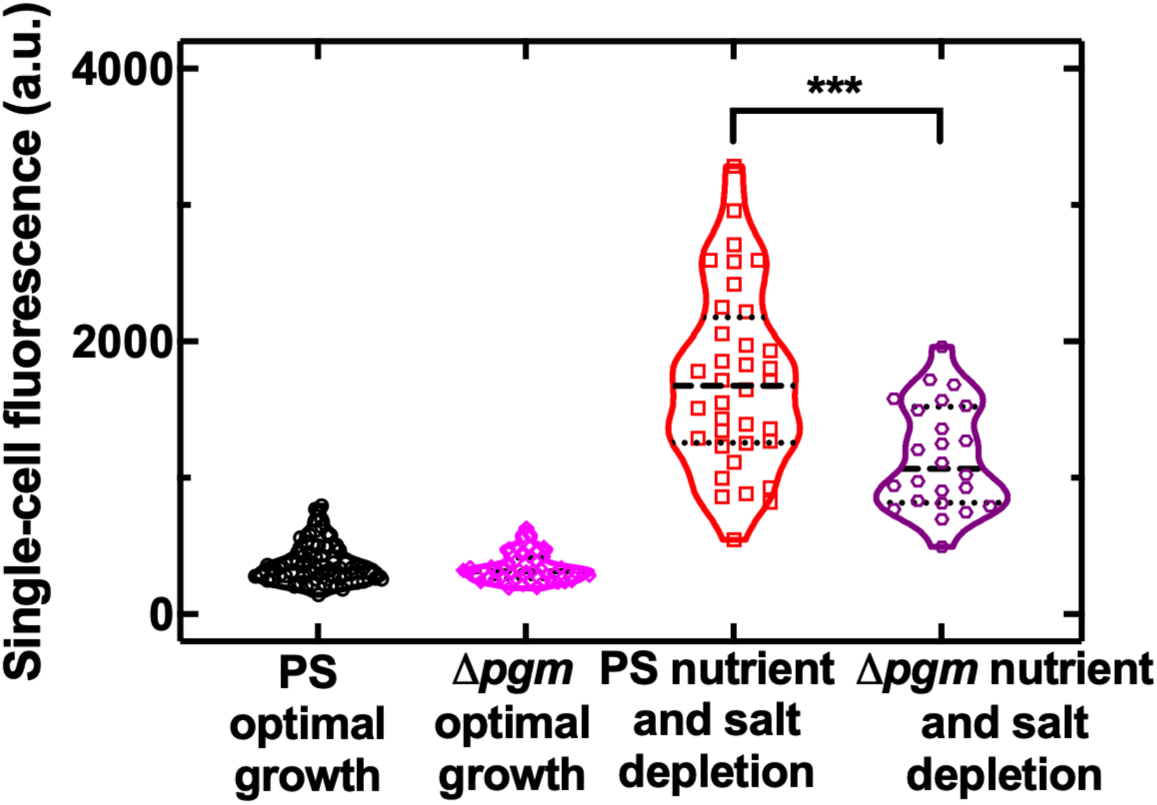
Distribution of single-cell fluorescence after 900 s incubation in 2-NBDG for the parental strain (PS) and Δ*pgm* deletion mutant under salt depletion or simultaneous nutrient and salt depletion. Dashed and dotted lines indicate the median and quartiles of each distribution, respectively. Under nutrient and salt depletion the Δ*pgm* deletion mutant displayed significantly lower 2-NBDG accumulation compared to the parental strain (***). In contrast, under optimal growth conditions the Δ*pgm* deletion mutant displayed 2-NBDG accumulation comparable to the parental strain.

**Figure S7.**
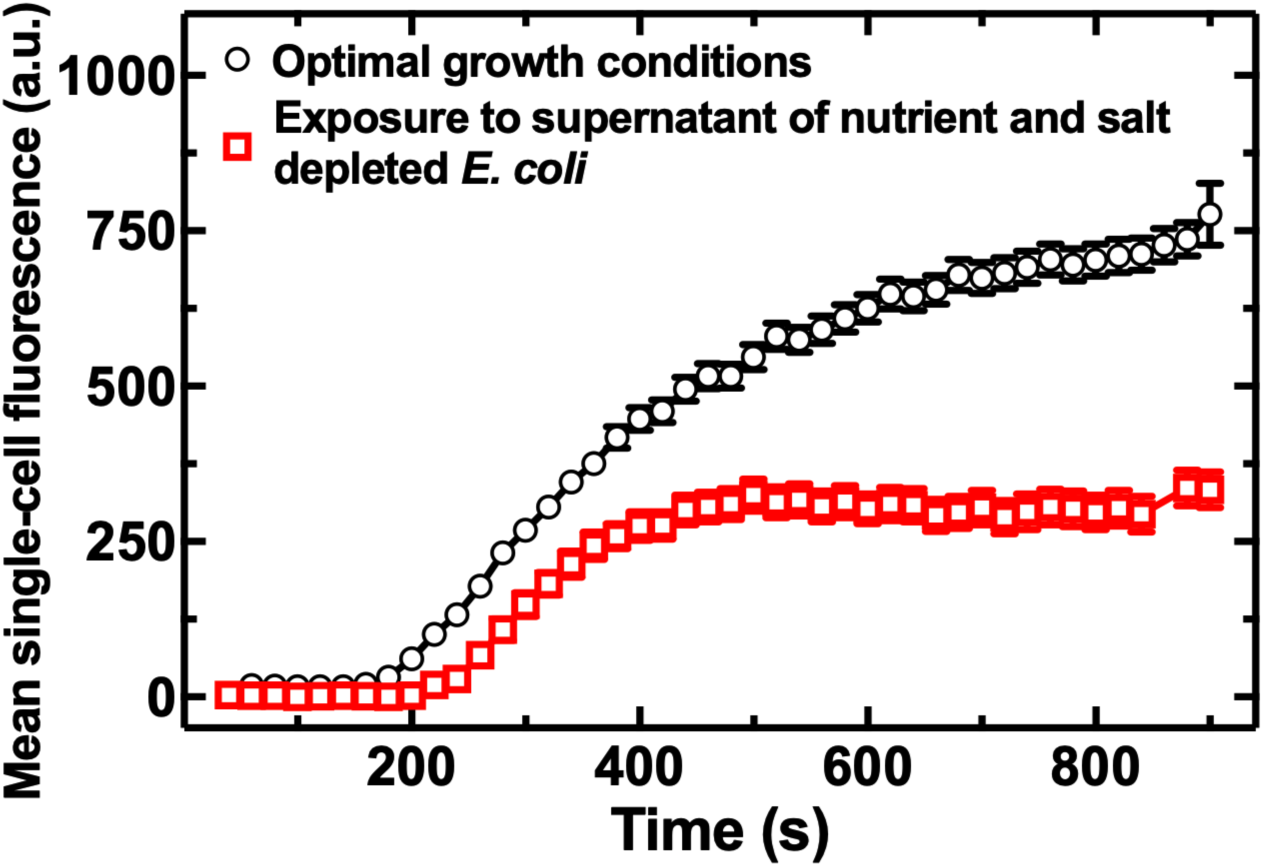
Temporal dependence of the mean intracellular fluorescence of the glucose analogue 2-NBDG in *E. coli* cultured in optimal growth conditions without (circles) or with an additional 1h exposure to the supernatant collected from *E. coli* cultures under combined nutritional and salinity depletion (squares) before 2-NBDG accumulation measurements. Symbols and error bars are the means and standard error of the means over at least 20 single-cell measurements.

